# A stable, hierarchical LIN code system for *Campylobacter jejuni* and *Campylobacter coli:* A unified genomic nomenclature for lineage-level typing and global surveillance

**DOI:** 10.64898/2026.02.10.705007

**Authors:** Kasia M. Parfitt, Ben Pascoe, Keith A. Jolley, Amy Douglas, Madison P. Goforth, Samuel K. Sheppard, Martin C.J. Maiden, Frances M. Colles

## Abstract

*Campylobacter* remains the leading cause of bacterial gastroenteritis worldwide, with *C. jejuni* accounting for around 90% of infection and *C. coli* accounting for most of the rest. Seven-locus multilocus sequence typing (MLST) has improved our understanding of host association and population structure, whilst core genome MLST (cgMLST), enables investigation of transmission events at high-resolution. However, the lack of a stable and standardised nomenclature for clustering of cgMLST data has limited reproducibility and long-term comparability between studies.

Here we introduce a joint, hierarchical Life Identification Number (LIN) code system that provides reproducible, multi-level genomic identifiers for *C. jejuni* and *C. coli* lineages. Using an updated cgMLST v2 scheme (1,142 loci) and globally representative datasets of high-quality genomes selected from over 53,000 assemblies in the *Campylobacter* PubMLST database (https://pubmlst.org/organisms/campylobacter-jejunicoli), we firstly defined LIN codes on a dataset of 5,664 genomes. Pairwise allelic distances were computed using MSTclust, and 18 nested thresholds were defined through silhouette, adjusted Wallace and adjusted Rand Index (ARI) statistics to capture the population structure from species to outbreak level resolution.

The LIN thresholds were then validated using a second dataset of 1,781 genomes from PubMLST and applied to a large water-associated outbreak dataset from New Zealand in 2016, containing clinical and ecological genomes. Further application of LIN codes was demonstrated by analyses of the *C. jejuni* ST-21 clonal complex and ST-6175 isolates, as well as the broader population structure of *C. coli,* using data from PubMLST. Across all datasets, LIN clusters were stable, largely monophyletic, and back-compatible with existing nomenclature, accurately distinguishing host-adapted and outbreak-associated lineages. By embedding cgMLST data within a stable and scalable nomenclature, the *Campylobacter* LIN system delivers consistent, automated genome-to-lineage assignment. This unified framework bridges population genetics and applied surveillance, enabling robust, real-time comparison of *Campylobacter* isolates across sources, studies, and time.

**Impact statement:** Human cases of *Campylobacter* worldwide continue unabated. Tracing the source of *Campylobacter* infection is particularly challenging given the sporadic or multi-source nature of outbreaks, with potential transmission from foodborne, animal or environmental sources. Seven-locus MLST has greatly improved our broad understanding of *Campylobacter* population structure. However, whilst high-resolution cgMLST alleles and STs themselves do not change, longitudinal cluster analyses of cgMLST data have lacked a stable nomenclature, rendering them unsuitable for robust and comparable surveillance over time. Life Identification Number (LIN) codes provide a solution to this problem, establishing an automated and scalable nomenclature derived directly from cgMLST profiles, that is stable over time. We have implemented a joint *C. jejuni* and *C. coli* LIN code scheme in PubMLST, with scripts for real-time lineage assignment. LIN codes are back-compatible with existing MLST nomenclature, and we demonstrate their added practical value for exploring population structure and high-resolution outbreak investigation. LIN codes support surveillance of *Campylobacter* in a One Health context, by enabling consistent typing at multiple levels across different sources, laboratories and time.

**Data summary:** 1. The isolate collections used to develop the LIN codes are publicly available and searchable as individual projects on the PubMLST database (https://pubmlst.org).

- **LIN code development (Dataset 1)** (n=5,664 isolates, up to 200 isolates per clonal complex)
- **LIN code validation (Dataset 2)** (n=1,781 isolates), up to 50 isolates per clonal complex
- **Outbreak investigation (Dataset 3)**: New Zealand 2016 Havelock North waterborne outbreak, Gilpin *et al* (n=161 isolates) [1]
- **Population structure exploration (clonal complex) (Dataset 4)**: ST-21 complex (n=1800 isolates, up to 100 isolates randomly selected from each country)
- **Population structure exploration (sequence type) (Dataset 5)**: ST-6175 (n=321 isolates, genomes with ‘good’ cgMLST v2 annotation)

2. The software for LIN code development is publicly available as follows:

- MSTclust for pairwise distance matrices https://gitlab.pasteur.fr/GIPhy/MSTclust [2]
- Python script to define LIN codes in a local dataset; (https://gitlab.pasteur.fr/BEBP/LINcoding)
- BIGSdb Perl script to define LIN codes from cgMLST profiles on the PubMLST database; (https://github.com/kjolley/BIGSdb/blob/develop/scripts/maintenance/lincodes.pl)

## Introduction

*Campylobacter* remains the leading bacterial causes of human gastroenteritis worldwide, with *Campylobacter jejuni* and *Campylobacter coli* responsible for an estimated 160 million global infections annually [3]. In the UK alone, *Campylobacter* accounts for >60,000 laboratory-confirmed cases each year, with a further increase of 17% reported in 2024 [4]. The burden extends beyond acute gastroenteritis, as *Campylobacter* is the predominant antecedent of Guillain–Barré (GBS) syndrome, contributing to up to 30% of GBS cases and often associated with its more severe form, [5]. Reproducible, high-resolution genomic tracking tools are essential to robustly identify transmission events, which may be complex for *Campylobacter* given the organism’s extensive reservoir amongst poultry, livestock, wild birds, companion animals, and environmental waters.

The public health importance of reliable genomic nomenclature is exemplified in outbreak investigations [1, 6–12]. The 2016 Havelock North waterborne outbreak in New Zealand is the largest campylobacteriosis outbreak recorded to date. There were between 6,260 and 8,320 people estimated to have been infected with *Campylobacter*, with 953 physician reported cases, three cases of GBS and at least four deaths attributable to the outbreak [1]. Whole-genome sequencing (WGS) of clinical isolates, as well as those from contaminated potable water and sheep faeces, revealed near-identical clusters of cgMLST genotypes linking animal, water, and human cases. This study demonstrated that high-resolution typing is essential to distinguish outbreak and background sporadic cases.

Multilocus sequence typing (MLST), using 7 loci, provides well-established nomenclature for *Campylobacter*, enabling global comparisons of sequence types (STs) and clonal complexes (CCs) (groups of closely related STs). In comparison, the higher discriminatory power of core genome MLST (cgMLST) results from indexing genetic variation at more than 1,000 loci[13]. Whilst cgMLST nomenclature such as allele and ST definitions are stable, single-linkage cluster definitions can change over time as new genomes are added to a dataset [14]. As a result, *Campylobacter* cgMLST clustering has lacked a consistent, stable and portable nomenclature analogous to MLST-defined clonal complexes.

To address these challenges, we introduce a stable, genome-wide nomenclature for *C*. *jejuni* and *C. coli* based on Life Identification Numbers (LINs). The LIN code concept was originally proposed for *Pseudomonas syringae* using Average Nucleotide Identity (ANI) analysis [15]. It has since been adapted for use with cgMLST data, which in comparison to ANI, gives greater reproducibility for distinguishing near-identical variants, and was first applied to *Klebsiella pneumoniae* [14]. The cgMLST based LIN approach has recently been extended to multiple other species, including *Klebsiella pneumoniae, Streptococcus pneumoniae, Staphylococcus aureus, Moraxella catarrhalis, Neisseria gonorrhoeae* and *Corynebacterium diptheriae* [14, 16–20]. LIN codes use a hierarchal clustering structure to group genomes, with each integer corresponding to predefined allelic divergence thresholds derived from cgMLST distances. Read from left to right, each position in the LIN code relates to increasing similarity between isolates. In other words, the degree of similarity between isolates can be assessed by the number of digits in the string (threshold number or ‘prefix’) at which they match, whilst those that diverge at the first threshold will be the most dissimilar.

In this study, we develop the first unified *C. jejuni* and *C. coli* LIN code nomenclature, built upon the updated and curated cgMLST v2 scheme comprising 1,142 loci. Due to the high levels of inter-species recombination, it was necessary to create both a cgMLST scheme and LIN codes that accommodate both *Campylobacter* species. Using a globally representative set of 5,664 high-quality genomes selected from >53,000 assemblies in PubMLST, we characterised allelic distance structure and defined 18 hierarchical thresholds using silhouette scores, adjusted Wallace coefficients and adjusted Rand indices. These thresholds were translated into a stable LIN code scheme and implemented directly into the PubMLST database where LIN codes can be assigned to new genomes in near real-time.

We validated the *Campylobacter* LIN codes across different epidemiological contexts, using a second globally representative validation dataset from PubMLST, and genome data from the 2016 Havelock North waterborne outbreak in New Zealand [1] (n=161 isolates were retrieved from public repositories). Further application of the LIN codes was tested using a globally representative dataset of ST-21CC and ST-6715 isolates from PubMLST (https://pubmlst.org)[21]. This work establishes a stable, scalable, and biologically meaningful whole-genome nomenclature for *Campylobacter*, designed to support global genomic epidemiology, outbreak investigation, and surveillance in a One Health context.

## Methods

### Development of a robust cgMLST v2 for *Campylobacter jejuni and Campylobacter coli* as the foundation of LIN codes

A robust cgMLST framework was required as the foundation for hierarchical LIN coding. We therefore re-evaluated the 1,343-locus cgMLST v1 scheme originally published in 2017 and assessed the performance of each locus across the current PubMLST *Campylobacter* collections [22]. All loci were screened for inconsistent start sites, internal stop codons, phase variation, paralogs, and frequent assembly or annotation failures. Loci exhibiting uncorrectable problems across >2% of publicly available genomes (threshold applied consistently to both *C. jejuni* and *C. coli*) were removed from the scheme. A total of 201 loci were excluded, yielding a revised cgMLST v2 scheme comprising 1,142 loci (Supplementary Table S1). Each retained locus was required to be present and reliably callable in ≥98% of high-quality genomes (≤100 contigs) from both species. The updated scheme was validated across all publicly available genomes with <=100 contigs (n=63,555), comparing allele call completeness and locus stability between v1 and v2. The v2 scheme is now the default cgMLST scheme used by PubMLST, while v1 remains available for backward compatibility.

### Globally diverse *Campylobacter jejuni* and *Campylobacter coli* genomes were used for LIN code development

All *C. jejuni* and *C. coli* genomes available in the PubMLST database on 25 January 2025 (n=86,715) were downloaded and assessed for quality. Species identity was confirmed using rMLST-based typing [23]. Genomes were retained if they contained ≤50 contigs and had ≥ 99% of cgMLST v2 loci assigned. A total of 41,659 genome assemblies met these criteria. To generate a representative dataset for LIN code development, a subset of 5,664 genomes (Dataset 1) were selected using a semi-randomised sampling procedure using the dplyr (v1.1.4) [24] function in RStudio (v4.5.2) [25], ensuring broad representation across clonal complexes, host sources, and geographic regions while limiting each CC to ≤50 isolates.

The resulting dataset (Dataset 1) comprised 5,224 *C. jejuni* and 440 *C. coli* genomes. Most clonal complexes (CCs) (n=42) associated with human disease, animal reservoirs, and environmental niches were captured. Three CCs, ST-1264CC, ST-1325CC and ST-1347CC, from wild birds and with 11 genomes or fewer were not included. Genome statistics such as assembly size, contig count, GC content, and N50 were downloaded from PubMLST and assessed to confirm data quality (Supplementary Figure 1a). The total number of contigs ranged from 1 to 50 (mean = 31, median = 32 contigs). The log₁₀ N50 ranged from 4.7 to 6.6 (mean = 5.3, median = 5.2), and the GC content (%) ranged from 29.88% to 34.41% (mean = 30.43%, median = 30.39%).

### Selection of LIN code thresholds

Using the LIN code development Dataset 1 (n=5,664), candidate LIN code thresholds were selected by visual analysis of discontinuity within the population structure (points of inflection or *local minima* on the ridgeline plots), combined with statistical analysis (Silhouette score and adjusted Wallace coefficient). The candidate LIN thresholds were then validated using adjusted Rand indices (ARI) and by comparison with existing MLST nomenclature to assess biological relevance (Datasets 1 and 2). Further validation of LIN code application was carried out using published[1] and investigative datasets drawn from PubMLST (Datasets 3-5).

In more detail, the first step for selecting LIN thresholds was to create a pairwise distance matrix for Dataset 1 genomes using MSTClust v0.21b (https://gitlab.pasteur.fr/GIPhy/MSTclust) [2] (Figure 1a). The output from MSTClust was manipulated in RStudio [25], using Reshape2 (v1.4.4) [26] to convert the matrices into long format. A histogram and ridgeline plots were drawn for the overall *Campylobacter spp.* population structure using the ggridges package from ggplot2 (v4.0.0) [27]. The ridgeline plots compared percentage allelic mismatches of the *C. jejuni* and *C. coli* isolates with: (i) Sequence Type (ST), (ii) ribosomal ST (rST), and (iii) clonal complex (CC) (Figure 1b). LIN code thresholds were assigned to points of discontinuity on the histogram and ridgeline plots, and the percentage (%) dissimilarity and corresponding number of allelic mismatches for each threshold noted.

**Figure 1.**
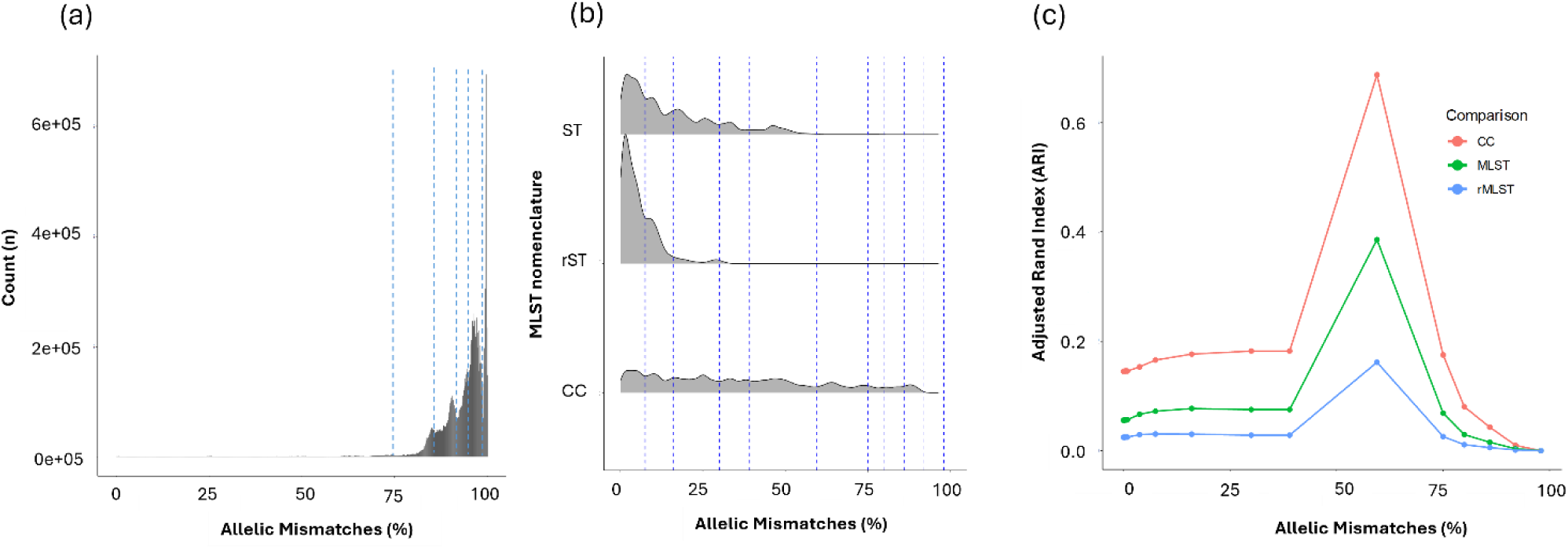
Plots used to identify distance thresholds for hierarchical cluster assignment; (a) pairwise allelic distance, (b) ridgeline plots of Dataset 1 matching the pairwise distance matrix to (i) clonal complex (CC), (ii) Sequence Type (ST), and (iii) ribosomal ST (rST) (key LIN thresholds are shown by dashed lines), (c) adjusted Rand index to assess the LIN code clustering and pre-defined metrics (ST, CC, and rST).

MSTClust v0.21b was used, with default parameters, to assess the Silhouette score [28] and adjusted Wallace coefficients [29] of the pairwise allelic mismatches of Dataset 1 [2, 20]. The Silhouette score was used to measure the cohesion of the clusters in the dataset. The adjusted Wallace coefficient indicated the robustness to subsampling bias within the cluster groups. A value of 1 indicated the best score (highest cohesion, most robust) for each of the tests respectively.

Once the 18 LIN thresholds were selected, a local LINcoding Python script (https://gitlab.pasteur.fr/BEBP/LINcoding) was used to assign draft LIN codes to each isolate in Dataset 1. This was used to check the suitability of the chosen thresholds and the accuracy of isolate assignment prior to PubMLST integration. The ARI [30] was calculated using the mclust package (11.0.4) and compared concordance of clustering between the designated LINs and CC, ST, and rST [31] (Figure 1c). An ARI value of 1 indicates perfect agreement between LINs and MLST classifications.

### Implementation and validation of LIN codes on PubMLST

The local LIN codes (https://gitlab.pasteur.fr/BEBP/LINcoding) from Dataset 1 were used to seed the PubMLST database [32]. After this, a minimum spanning tree was used to order the isolates for automated LIN code assignment using PubMLST, run in batches of 10,000 isolates [14]. LIN codes are automatically assigned on PubMLST for newly uploaded genomes with a core gene Sequence Type (cgST) on a weekly basis [20]. For phylogenetic validation of the LIN codes following implementation on PubMLST, it was necessary to use a smaller dataset due to practical limitations when using tools such as the Genome Comparator plugin on PubMLST [21]. The smaller dataset (Dataset 2) consisted of 1,781*Campylobacter* genomes (n=1,663 *C. jejuni* and n=118 *C. coli*) from up to 50 isolates for each clonal complex. The dataset was globally diverse, representing six continents, 41 countries, and 42 clonal complexes from a range of sources. High quality genomes were selected, and statistics such as genome assembly size, contig count, GC content and N50 (Supplementary Figure 1b).

Nucleotide alignments of the cgMLST loci (n=1,142) for the genomes in Dataset 2 were created using the Genome Comparator plugin on PubMLST. A cgMLST MUSCLE alignment with ‘all loci’ was selected, with all other parameters using default settings. A maximum likelihood phylogenetic tree was created from the core genome alignment output using FastTree (v2.1.11) [33]. The FastTree output was then subject to ClonalFrameML (v1.12) [34] to adjust for recombination events (Figure 2). Association of LIN code prefixes with MLST-defined CCs for Dataset 2 was further validated across all *C. jejuni* and *C. coli* isolates on the PubMLST database (Supplementary table S2). Unique LIN prefixes (minimum defining series of integers based on cgMLST) were subsequently implemented in PubMLST as ‘nicknames’ (eg ST-21 clonal complex (cgMLST v2)) and are searchable for each of the MLST-defined CCs[14].

**Figure 2.**
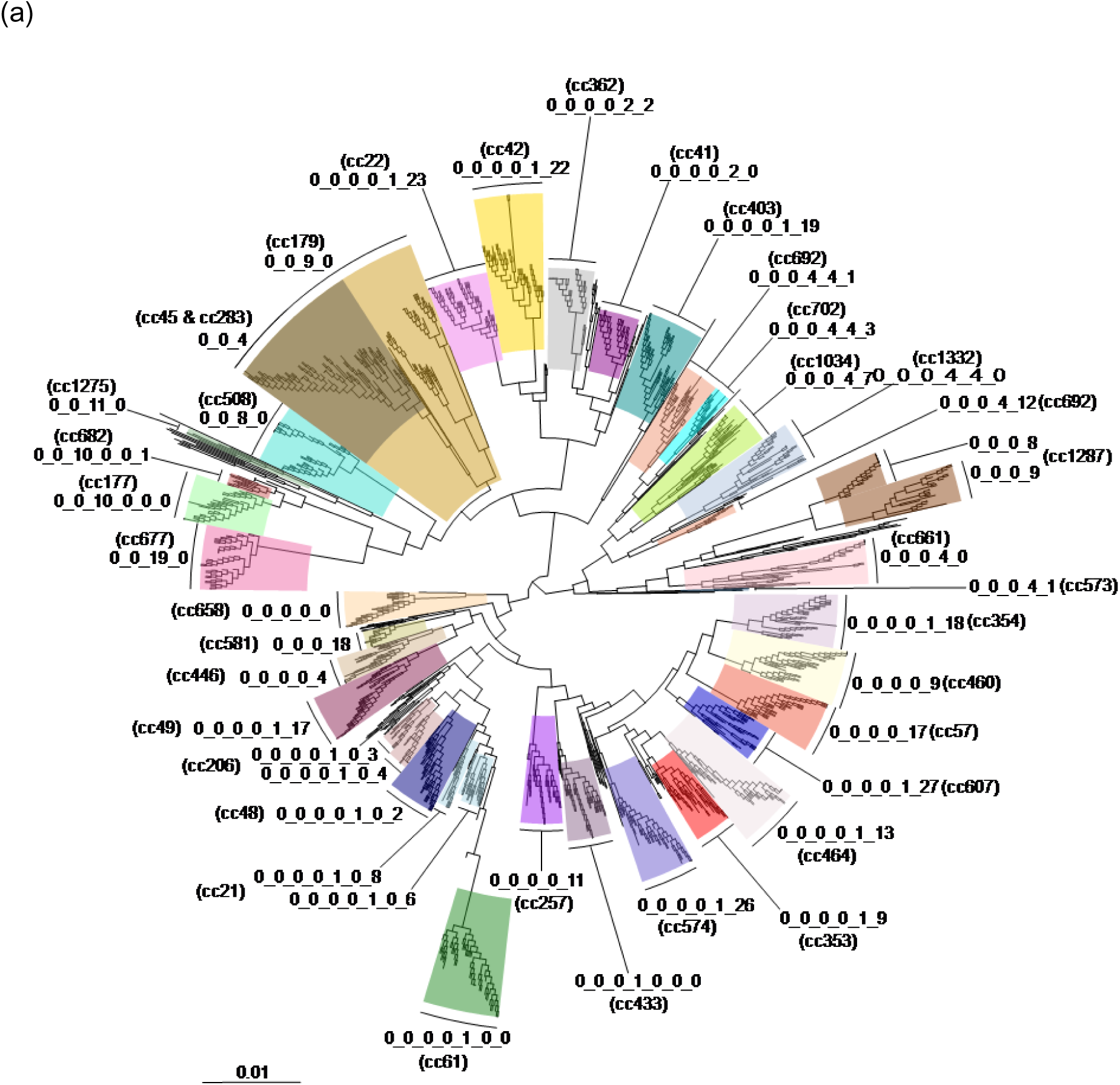
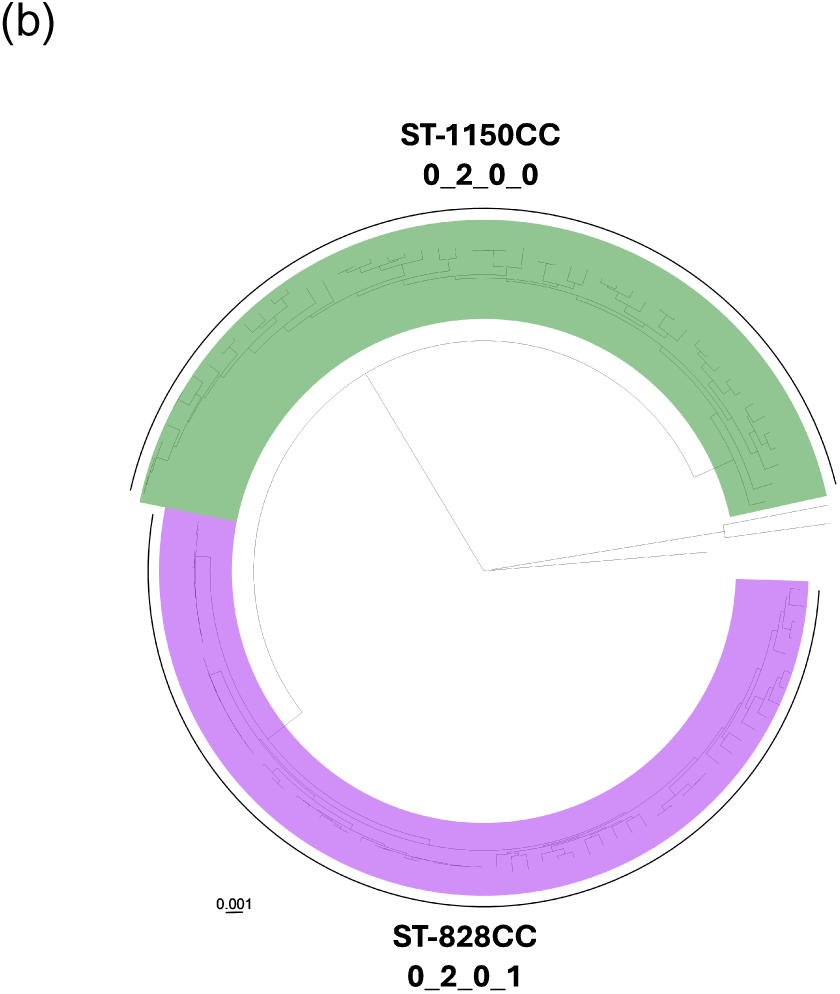
Validation of LIN code thresholds (a) *C. jejuni* and (b) *C. coli* from Dataset 2, globally representative isolates from Dataset 2. Clonal complexes are indicated by different colour blocks, with LIN codes shown to distinguishing thresholds.

### Example application of *Campylobacter* LIN codes to clinical and environmental datasets

Application of the LIN codes to an outbreak setting was tested using the previously published 2016 Havelock North waterborne outbreak dataset from New Zealand [1]. The dataset used in this study comprised 161 isolates, of which 143 were from human stools, 6 from potable water and 12 from sheep faeces (Dataset 3). Most isolates were *C. jejuni*, with three *C. coli* isolates from human stool samples. Isolates grouped into nine clonal complexes, including most of the ST-42CC (n=58), ST-21CC (n=53) and ST-45CC (n=20), demonstrating a mixed source outbreak. Genome alignments and phylogenetic trees for these datasets were created and adjusted as above.

The established LIN codes were used then to investigate ST-21CC, which has proven problematic for source attribution analyses due to multi-source association [35]. We used a dataset of high-quality genomes, randomly selected for up to 100 isolates per country (Dataset 4). A total of 1,800 isolates spanning the years 1995-2024 encompassed six continents (excluding Antarctica), 43 countries, and a range of sources (human disease (n=1295, 71.9%), poultry (n=293, 16.3%), ruminants (n=156, 8.7%) and other (less than 35 isolates, 1.9% per source). There were 143 STs, of which ST-50 (n=454, 25.2%), ST-21 (n=252, 15.7%) and ST-19 (n=226, 12.6%) were the most common. The remaining STs accounted for less than 117 isolates (6.5%) each.

Within ST-21CC, it was chosen to examine ST-6175 in greater detail due to its prominence and frequent co-resistance to fluoroquinolone and tetracycline in a recent UK study [36]. We used LIN codes to investigate global differences in a dataset of 321 high-quality ST-6175 genomes selected from PubMLST (Dataset 5, accessed on 6^th^ October 2025). The earliest recorded ST-6175 was from a UK free-range broiler breeder chicken in 2004, before reaching a peak of 95 isolates recorded in 2021. Isolates in this dataset are recorded from 16 countries across four continents (Europe (n=236, 73.5%, North America (n=27, 19.3%), Asia (n=7, 2.2%), Africa (n=1, 0.3%) and unrecorded (n=15, 4.7%), with the majority coming from the UK (n=195, 60.7%) and the US (n=62, 19.3%).

Clustering of genomes within the ST-21CC and ST-6175 were compared with LIN codes and metadata such as continent, year, source and AMR profile and visualised using the PubMLST GrapeTree plugin [37]. The cgMLST v2 scheme and MSTree V2 options were selected. GrapeTree created a minimum spanning tree using the cgMLST allelic profiles, and clusters those that share the most alleles [20].

## Results

### The updated cgMLST (v2) can be applied more widely for *Campylobacter jejuni* and Campylobacter coli

The cgMLST scheme was updated to v2 which consists of 1,142 loci, removing 201 problematic loci from v1. These included hypothetical proteins, loci with putative or unknown function and pseudogenes (165/201, 82.1%), loci associated with metabolism (19/201, 9.5%), transportation (7/201, 3.5%) structural proteins (2/201, 1%), toxin/virulence (2/201, 1%), 50S ribosome (1/201, 0.5%), DNA replication (1/201, 0.5%), transcription factor (1/201, 0.5%), signal transduction (1/201, 0.5%) genetic information processing (1/201, 0.5%) and a chromosome associated protein (1/201, 0.5%) (Supplementary table S1). The threshold of missing loci allowed for isolates has been reduced from 50 loci in version 1 [22] to 25 or fewer missing loci in version 2. By implementing the cgMLST v2 scheme on PubMLST, cgSTs and clustering algorithms could be applied to 98.4% of *C. jejuni* isolates and 99.5% of *C. coli* isolates with genome assemblies containing 100 or fewer contigs using automated scripts (accessed on 13^th^ January 2026). This represents an improvement over the number of isolates with 100 or fewer contigs that were assigned cgSTs using the cgMLST v1 scheme (96.4% of *C. jejuni* and 95.6% of *C. coli* isolates).

### Distance structure defined by cgMLST gave 18 LIN thresholds

Using Dataset 1, 18 thresholds (ranging from 0 to 1,119 allelic differences) were chosen to match discontinuity in the population structure. These were shown by uneven distribution of pairwise allelic differences on the histogram, and discontinuity on the ridgeline plots (Figure 1). The final LIN threshold selection was supported by statistical analysis to maximise cluster cohesion and stability. The LIN thresholds were also validated by comparison with existing MLST nomenclature to assess biological relevance (Figure 1, Table 1). The pairwise allelic distance distribution exhibited clear multimodality and a negative skew on the histogram, consistent with other bacterial species populations [2, 17, 18, 20] (Figure 1a). The most common frequency of pairwise allelic mismatches peaked at around 80% similarity, which is consistent with there being two species, *C. jejuni* and *C. coli*[38]. The greatest difference of 1,142 allelic mismatches, which is all the core genes in the scheme, demonstrates high genetic diversity amongst the isolates, which is captured at the first LIN threshold.

**Table 1.** LIN thresholds, showing percent identify and allelic mismatch, and rank*, enabling back-compatibility with existing nomenclature. *Rank is an approximation of existing nomenclature and not definitive, for example, there is some overlap between CC and ST at LIN threshold-6 in practice.

| Threshold | 1 | 2 | 3 | 4 | 5 | 6 | 7 | 8 | 9 |
| --- | --- | --- | --- | --- | --- | --- | --- | --- | --- |
| % dissimilarity | 97.99 | 95.01 | 85.99 | 80 | 75.04 | 59.54 | 38.97 | 30 | 16.02 |
| % identity | 2.01 | 4.99 | 14.01 | 20 | 24.96 | 40.46 | 61.03 | 70 | 83.98 |
| Allele mismatch | 1119 | 1085 | 982 | 914 | 857 | 679 | 445 | 343 | 183 |
| Allele identity | 23 | 57 | 160 | 225 | 285 | 463 | 697 | 799 | 959 |
| Rank (LIN) | Genus | Introgressed species |  | Lineage (CCs) |  |  | Sub-lineage (nested CCs, STs) |  |  |

| Threshold | 10 | 11 | 12 | 13 | 14 | 15 | 16 | 17 | 18 |
| --- | --- | --- | --- | --- | --- | --- | --- | --- | --- |
| % dissimilarity | 7.5 | 3.77 | 0.9 | 0.61 | 0.44 | 0.26 | 0.18 | 0.09 | 0 |
| % identity | 92.5 | 96.26 | 99.1 | 99.39 | 99.56 | 99.74 | 99.82 | 99.91 | 100 |
| Allele mismatch | 86 | 43 | 10 | 7 | 5 | 3 | 2 | 1 | 0 |
| Allele identity | 1056 | 1099 | 1132 | 1135 | 1137 | 1139 | 1140 | 1141 | 1142 |
| Rank (LIN) |  |  | Clonal groups |  |  |  |  |  | Indistinguishable |

A high-level boundary of 1,119 locus differences (97.99% allelic mismatches) was chosen to define the ‘genus’ threshold, encompassing all *C. jejuni* and *C. coli* isolates including ‘ancestral’ *C. coli* lineages from clades 2 and 3, but distinguishing some small clusters of more distantly related environmental isolates (Table 1) [38].

A threshold boundary of 1,085 locus differences (95.01%) waschosen to define the ‘introgressed’ *C. jejuni* and *C. coli* ST-828 and ST-1150 CCs that are predominantly isolated from clinical and livestock isolates. These thresholds correspond to cut-off values between 0.95 and 0.99, which had normalised area under the curve (nAUC) Silhouette scores of 1.00.

The threshold boundaries of 857 (75.04% allelic mismatches) and 680 (59.54% allelic mismatches) represented most CCs and were defined as ‘lineage’, which encompasses the optimal threshold for lineage assignment based on the nAUC Silhouette and adjusted nAUC Wallace scores of 1.0 and 0.9, respectively. Furthermore, the 75.04% allelic mismatch threshold corresponded to an nAUC Silhouette score of 1.0 and an adjusted nAUC Wallace score of 0.78. Cut-off values between 59.19% and 59.86% had nAUC Silhouette scores of 0.99, and adjusted nAUC Wallace scores between 0.82-0.84. A ‘sub-lineage’ threshold was defined at 445 locus differences (38.97% allelic mismatches), selected for more distantly related or nested CCs. This corresponded to a cut-off value of 38.95%, with an nAUC Silhouette score of 0.95 and adjusted nAUC Wallace score of 0.94. The sub-lineage threshold for nested CCs overlaps with STs (range thresholds 7-9, 445 – 183 allelic mismatches).

An additional six threshold boundaries were included for epidemiological analysis and high-resolution investigation, defined as ‘clonal groups’. These correspond to 10, 7, 5, 3, 2, and 1 locus difference(s) (corresponding to 0.88%, 0.61%, 0.44%, 0.26%, 0.18%, and 0.09% allelic mismatches, respectively). This is consistent with boundaries defined in other published LIN code papers [2, 16, 20]. Isolates with indistinguishable LIN codes were identified through a 0-locus difference boundary (0% allelic mismatches).

### LIN thresholds accurately capture *C. jejuni* and *C. coli* species structure

Using Dataset 2, agreement of LIN code clusters with established MLST classification (ST, CC and rST) was assessed using the ARI. The 59.5% allelic mismatch threshold (680 locus differences) had the greatest concordance with all three metrics with an ARI score of 0.39, 0.69, and 0.16 for ST, CC, and rST, respectively (Figure 2c). The other thresholds had an ARI of between 0 and 0.18. These results illustrate a lower concordance between the ARI metrics and the designated LIN codes compared to schemes for other bacterial species[2, 16, 20]. This is likely attributable to (i) the mixed *C. coli* and *C. jejuni* population and (ii) the imperfect assignment of some CCs in the existing 7-locus scheme. With these complications, the ability of LIN codes to provide multilevel prefixes gives a better solution in defining the population structure within *C. jejuni* and *C. coli* and assessing clustering levels (Figure 2, and Supplementary table S2). Examples of imperfect matches between CC and LIN threshold which would affect the ARI metrics include the following; ST-45CC and ST-283C, indistinguishable at the 3-integer LIN prefix; ST-1287CC which has two LIN codes at the 4-integer LIN prefix; and ST-21CC which can only be distinguished from closely related CCs ST-48CC and ST-206CC with a 7-integer LIN prefix. The different prefix lengths give insight into these relationships with greater detail than MLST. Whilst the Silhouette and adjusted Wallace scores demonstrated cohesive and robust LIN thresholds, it was not possible to resolve low ARI scores and therefore the performance of LIN codes matching existing MLST nomenclature was instead validated across the whole PubMLST database, with emphasis on biological interpretation and robust functionality for investigative applications. Results are given as a range of matching thresholds rather than a single threshold, as befits the *Campylobacter* population structure.

Overall, there were 70,235 unique LIN codes assigned to 90,329 *C. jejuni* and *C. coli* isolates on PubMLST (22^nd^ December 2025). There were five LIN codes assigned to the ‘genus’ threshold. The predominant LIN code at this threshold was ‘0’, with 90,024 (99.7%) associated isolates. The remaining isolates were assigned to LIN code prefixes of ‘1’, ‘2’, ‘3’, and ‘4’, harbouring 18, 264, 15, and 8 isolates, respectively. These prefixes contained isolates that were unassigned to any clonal complex, and most were associated with wild bird and environmental water isolates. *C. jejuni* isolates were captured by prefixes 1, 3 and 4, with prefix ‘1’ related to *C. jejuni* isolates from wild birds in the US, and prefixes ‘3’ and ‘4’ related to distinct clusters of *C. jejuni* isolates containing both from wild birds and environmental waters from Europe. Prefix ‘2’ related to *C. coli* isolates distributed across five continents, with a large proportion from environmental waters (110/264, 41.7%), with human disease the next most common source (75/264, 28.4%).

There were 16 unique LIN codes at the 2^nd^ threshold, known as the ‘introgression’ group. Two of these (0_0, and 0_2) represented most (89,680/90,329 isolates, 99.3%) of the population. LIN prefix 0_0 (63,001/90,329 isolates,69.7%) was associated with *C. jejuni* and LIN code 0_2 (26,679/90,329 isolates, 29.5%) was associated with *C. coli* ST-828CC and ST-1150CC. LIN prefixes 0_3 to 0_7 contained between 1 and 88 isolates, all unassigned to CCs. Isolates with LIN prefix 0_4 were globally distributed *C. jejuni* and notably associated from cattle sources. Isolates with LIN prefix 0_6 were *C. coli,* also globally distributed and were predominantly associated with wild bird and environmental water sources.

### LIN thresholds are phylogenetically coherent and compatible with existing MLST nomenclature

To validate the biological integrity of LIN clusters, we constructed two recombination-adjusted maximum likelihood phylogenies using representative subsets of 1,663 *C. jejuni* (Figure 2a) and 118 *C. coli* (Figure 2b) genomes from Dataset 2. LIN-defined groups were largely congruent with monophyletic clades in the tree for both species, indicating that the cgMLST-based thresholds capture evolutionary structure even in the presence of frequent recombination. The results were further validated with all genomes with cgST data on PubMLST (22^nd^ December 2025).

At mid-range thresholds (e.g. LIN thresholds 5–7, which represent ‘lineage’ and ‘sub-lineage’), clusters aligned closely with CCs and previously defined host-associated ecotypes (Figure 2, Table 1, Supplementary Table S2). Some CCs resolved a little sooner. For example, ST-45CC and ST-283CC shared the same LIN prefix 0_0_4 (Figure 2, Supplementary Table S2, Supplementary Figure S2) and group closely with ST-179CC with LIN prefix 0_0_9. It was not until the 10^th^ LIN threshold that ST-283CC (0_0_4_0_0_0_0_0_14_1) could be distinguished from ST-45CC, suggesting that the originally described ST-283CC is similar to an ST level of resolution. ST-45CC is recognised as a host ‘generalist’ [35] and is often isolated from environmental sources, uniquely peaking in the summer in UK human disease [39]. These characteristics are consistent with the early distinguishing LIN prefix and closest grouping with ST-179CC associated with wild birds on the FastTree (Figure 2, Supplementary Figure S2).

Seven CCs (ST-1034CC, ST-1332CC, ST-283CC, ST-573CC, ST-661CC, ST-692CC, and ST-702CC) that shared the same LIN prefix at the 4^th^ threshold (0_0_0_4), which were resolved at the 5^th^ LIN threshold. Notably, the CCs ST-692CC, ST-702CC, ST-1034CC and ST-1332CC are associated with wild birds, particularly geese[40, 41] and group together on the FastTree (Figure 2). The CCs ST-573CC and ST-661CC group nearby on the FastTree are associated with chicken. There were 6 CCs (ST-21CC, ST-179CC, ST-508CC, ST-581CC, ST-677CC and ST-1275CC) that resolved at the 5^th^ LIN threshold. The host ‘generalist’ ST-21CC is described in greater detail below. At the 6^th^, 7th, and 9^th^ thresholds, 8, 16, 5, and 1 CC resolved respectively. No CCs resolved at the 8^th^ threshold. ST-1287CC was associated with two prefixes at the 4^th^ LIN threshold (0_0_0_8 and 0_0_0_9), suggesting it could potentially have been assigned to two different CCs using the original MLST scheme for consistency in the level of resolution. The LIN 0_0_0_8 prefix had a subcluster from ‘other animals’, which could be the reason they are distinct from the 0_0_0_9 prefix which was more common amongst wild birds.

It has been previously noted that ST-353CC is polyphyletic [42]. Analysis shows most ST-353CC isolates grouped in the LIN 0_0_0_0_1_9 prefix (Supplementary table S2). Of 7,929 isolates with this prefix on PubMLST (22^nd^ December 2025), 7721 (97.6%) grouped into ST-353CC. However, ST-353CC isolates could be found in a further 10 prefixes at the 6^th^ LIN threshold, with the next most common being 0_0_0_0_1_5 (135/353 ST-353CC isolates, 41.3%) and 0_0_0_0_1_8 (114/138 ST-353CC isolates, 52.6%).

### Application to outbreak and population datasets

#### Isolates with identical LIN code identified amongst the 2016 Havelock North waterborne outbreak, New Zealand

We applied the LIN code system to a previously published waterborne outbreak dataset from New Zealand to assess its epidemiological resolution [1]. Our results using LIN codes identified the same clusters of identical clonal groups, linking human clinical isolates with those from sheep and potable water (Figure 3). Reanalysis of the New Zealand outbreak dataset[1], using the updated cgMLST scheme, identified 20 fewer cgSTs (140 cgSTs in cgMLST v1 compared to 120 cgSTs using the updated cgMLST v2 scheme). A total of 109 LIN codes were identified amongst the 161 New Zealand outbreak isolates, of which, three LIN codes accounted for 9 to 19 isolates respectively, seven LIN codes accounted for 2-5 isolates respectively, with the remaining 99 LIN codes were each recovered only once.

**Figure 3.**
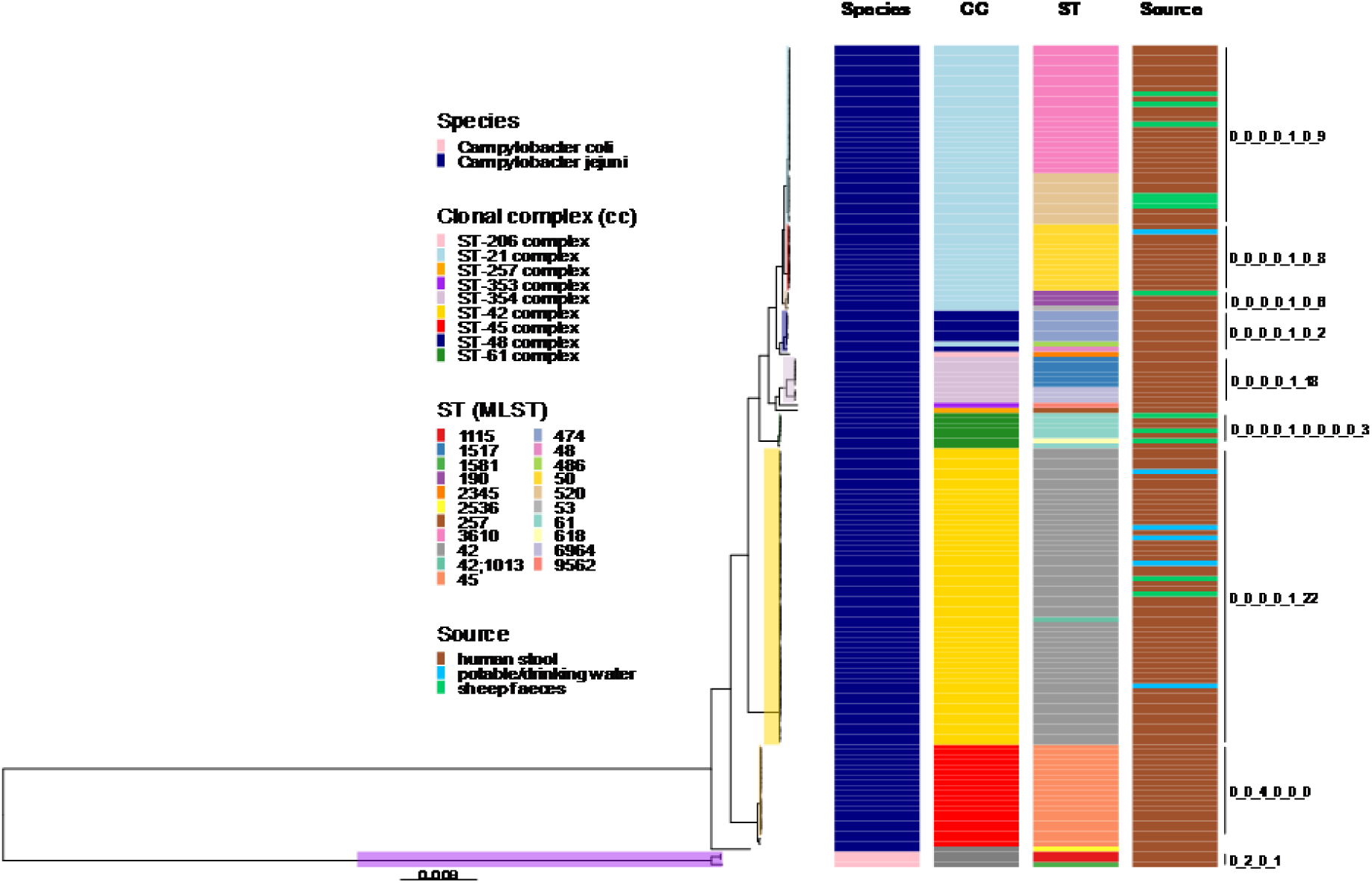
Phylogenetic tree showing *Campylobacter* isolates from the waterborne Havelock North outbreak in New Zealand, 2016. Each LIN code grouping is highlighted by different colour blocks, matched to clonal complex (CC).

The advantage of using LIN code nomenclature over cgSTs was that isolates matching at all 1,142 cgMLST loci were immediately identifiable by their matching 18-digit LIN code. Isolates from either potable water or sheep faeces were found to have identical matching LIN codes with clinical isolates on four occasions; however, no single LIN code was found to match across all three sources. The most common match was for isolates typed as ST-42 (ST-42 complex) with LIN code 0_0_0_0_1_22_0_0_7_1_1_1_0_0_0_0_0_0, accounting for 17 isolates from human disease and two isolates from potable water (Figure 4b). A second, smaller cluster of five ST-42 isolates with a matching LIN code 0_0_0_0_1_22_0_0_7_1_1_1_0_0_0_0_0_3 were isolated from human disease (n=4) and potable water (n=1). A further 26 LIN codes were associated with ST-42 isolates from the outbreak, most found only once, but one identified nine times from human disease.

**Figure 4.**
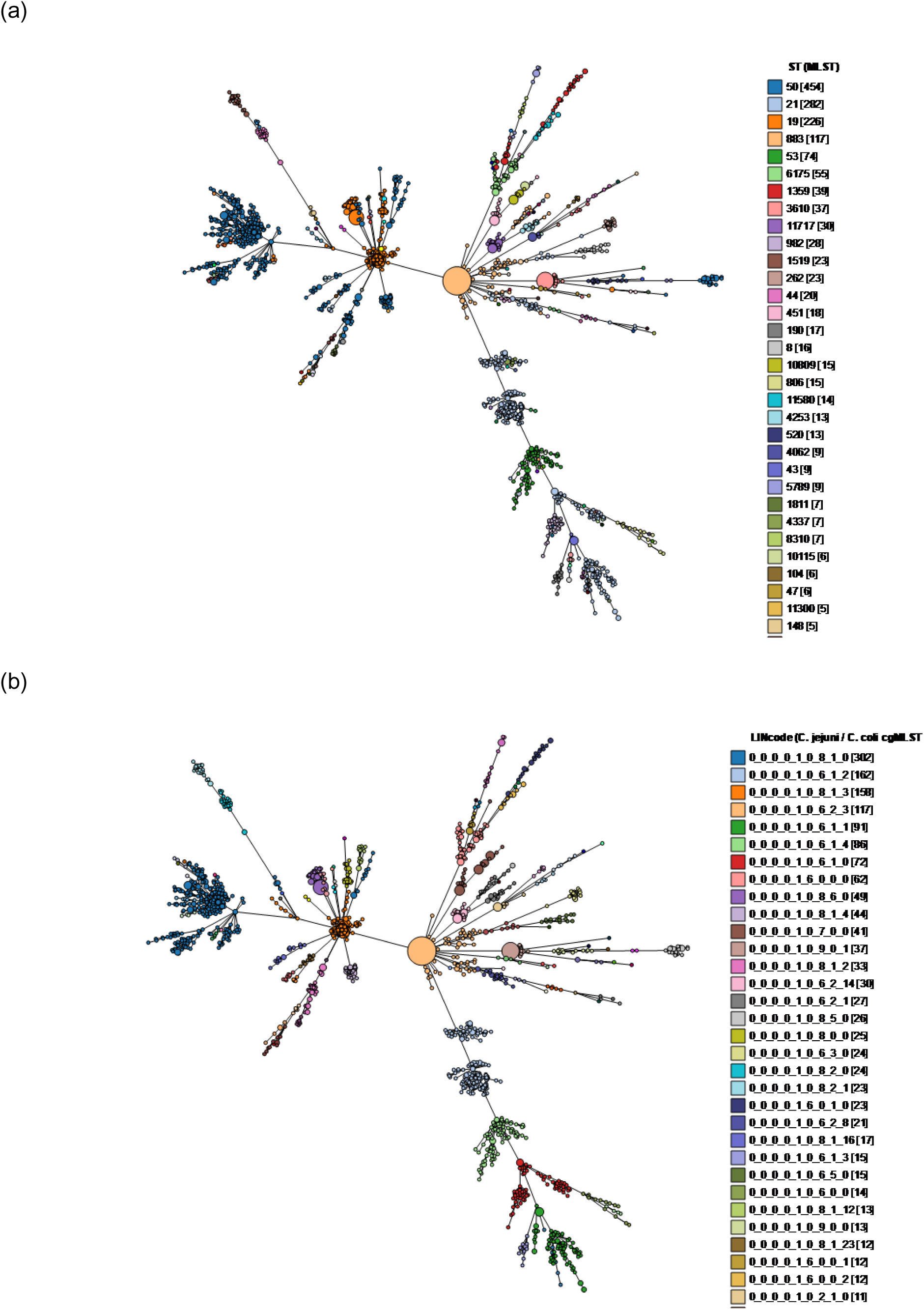
The clustering of isolates within the ST-21CC test dataset using the cgMLST v2 scheme and labelled by (a) ST and (b) LIN code at 9th threshold, shown using GrapeTree.

Isolates with a LIN code matching from human disease and sheep sources from the same New Zealand waterborne outbreak were identified amongst two STs grouping into ST-21 complex, ST-3610 and ST-190. A large cluster of 15 ST-3610 isolates with matching LIN code 0_0_0_0_1_0_9_0_1_0_0_0_0_0_0_0_0_0 (Figure 4) were identified amongst 13 human disease isolates and two isolates from sheep. A further ten ST-3610 isolates were identified with LIN codes seen only once, accounting for nine human disease isolates and two sheep isolates. ST-190 isolates were identified three times in the outbreak, with one human disease and one sheep isolate having a matching LIN code, and a third human disease isolate differing at two loci.

#### LIN codes resolve sub-clustering within the *C. jejuni* generalist ST-21 clonal complex

All 1,800 isolates from the test ST-21 clonal complex dataset were encompassed by the same LIN code at the 5^th^ threshold (0_0_0_0_1) which is an equivalent threshold to other *Campylobacter* CCs. Within the ST-21CC, the 9^th^ LIN threshold gave an equivalent level of resolution as 7-locus MLST, with 143 STs and 98 LIN codes identified at the 9^th^ threshold (Figure 4). The distribution was a little more even using the LIN codes at the 9^th^ threshold, with the most common ST-50 being identified 454 times, compared to the most common LIN code 0_0_0_0_1_0_8_1_0 which was identified 302 times (Supplementary figure S3).

However, some STs within the ST-21CC were distinct at a lower threshold, with the 6^th^ LIN threshold encompassing most isolates from the most three most common STs, ST-50 (n=454), ST-21 (n=282) and ST-19 (n=226). There was a prevalent sub-lineage within each ST, but the 12^th^ LIN threshold (STs 50 and 21) and the 15^th^ LIN threshold (ST-19) were needed to identify small sub-clusters (‘clonal groups’) of 20-30 isolates within these STs.

#### LIN codes reveal antimicrobial resistance associated sub-structure within ST-6175 (ST-21CC)

Each of the 321 ST-6175 isolates from the test dataset were encompassed by the 9^th^ LIN threshold, equivalent to the threshold range for STs in the other datasets we tested. Within ST-6175 however, isolates were found to cluster into clonal groups at the 11^th^ LIN threshold, which matched continent, predicted resistance pattern to fluoroquinolone and tetracycline resistance, and resistance allele using GrapeTree analysis (Figure 5 and Supplementary Figure S4). The clonal groups could be distinguished as follows; North American (LIN 0_0_0_0_1_6_0_0_0_0_0, fluoroquinolone resistance with CAMP0950 allele 1537, tetracycline sensitive), European (LIN 0_0_0_0_1_6_0_0_0_0_2, fluoroquinolone resistance with CAMP0950 allele 176, tetracycline resistance with CAMP1698 allele 13), Asian (LIN 0_0_0_0_1_6_0_0_0_0_4, fluoroquinolone resistance with CAMP0950 allele 61, tetracycline resistance with CAMP1698 allele 46), and one African isolate (LIN 0_0_0_0_1_6_0_0_0_4_0, fluoroquinolone resistance with CAMP0950 allele 1498, tetracycline sensitive). These findings are consistent across time. All except one of the ST-6175 isolates had the T86I *gyrA* mutation which is highly predictive of fluoroquinolone resistance. The exception was a single cattle isolate from Luxembourg in 2017, which clusters with other resistant isolates in the European clonal group, matching at the ninth threshold LIN code.

**Figure 5.**
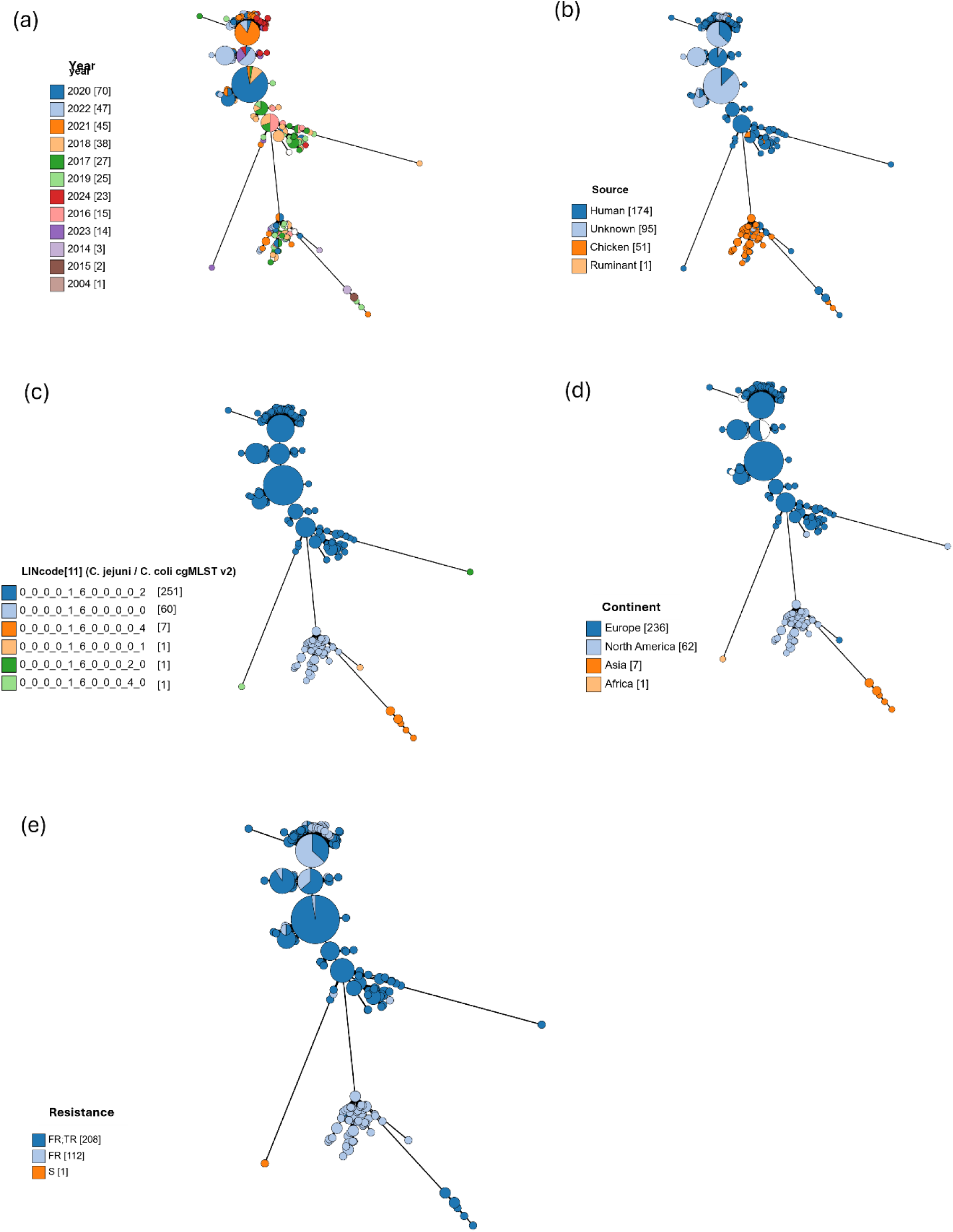
The clustering of isolates within ST-6175 (ST-21CC) using the cgMLST v2 scheme and labelled by (a) year, (b) source, (c) 11^th^ LIN threshold, (d) continent and (e) antimicrobial resistance profile.

#### LIN codes give greater insight into the population biology of *C. coli* and are more accurate than STs

In contrast to the 45 CCs identified to date by 7-locus MLST within *C. jejuni*, there are just two CCs identified within *C. coli* (ST-828CC and ST-1150CC), together with two ancestral clades with divergent lineages[38]. In test Dataset 1, all genomes within the large, multi-host ST-828CC were identified at the fourth threshold LIN code 0_2_0_1 and *vice versa* (Figure 2b, Supplementary Figure S5). Extending the analysis to all good quality ST-828CC genomes assigned a LIN code and with 100 or fewer contigs on PubMLST, 99.9% (19,136/19,147) had the prefix of 0_2_0_1 (accessed 1^st^ February 2026). The exceptions included ten ST-828CC isolates from pig and beef sources in Peru with 4-threshold LIN code 0_2_0_1, and one US chicken isolate with 4-threshold LIN code 0_2_0_0. This isolate contained four MLST loci in common with ST-828 and three MLST loci in common with ST-1150. All ST-1150CC genomes in Dataset 1 were identified by LIN code 0_2_0_0 at threshold 4 in all isolates. This same 4-threshold code also accounted for all 443 isolates on PubMLST, for which a LIN code was assigned. Isolates from the *C. coli* ancestral clades could be distinguished by LIN codes at the second threshold, for example, 0_3 and 0_6.

As with *C. jejuni,* resolution of clusters at approximately the same level of ST within the *C. coli* population occurred around the seventh LIN threshold (Supplementary Figure S5). Within the large multi-host ST-828CC however, some of the more prevalent STs such as ST-828, ST-829 and ST-854 began to split at the 6^th^ LIN threshold. Greater resolution was achieved through the increasing LIN thresholds, with 22/262 STs splitting at the 6^th^ LIN threshold, 29/262 STs splitting at the 7^th^ LIN threshold and 36/292 STs splitting at the 8^th^ LIN threshold. Comparing the clustering using GrapeTree (Supplementary Figure S5) and cgSTs, the LIN thresholds, unsurprisingly, give better representation of sub-clusters within ST-828CC than STs based upon 7-locus MLST.

Within ST-828, a prevalent ST within the ST-828CC, there is coherent sub-clustering shown at the 6^th^ LIN threshold (Supplementary Figure S6). The clusters are, in the main, divided by continent (Europe *vs* North America and Asia) and source (poultry *vs* pigs and ruminants).

## Discussion

In this study, we present a stable, genome-wide clustering nomenclature for *Campylobacter jejuni* and *Campylobacter coli* based on Life Identification Numbers (LINs) that offers an automated and scalable enhancement to whole genome-based sequence analyses. Typing of these two *Campylobacter* species can be problematic given the high levels of recombination within and between species, and complex transmission pathways involving a wide range of isolation sources. The newly presented and validated LIN codes offer a nested, string-based nomenclature where each integer encodes a cluster of genetic relatedness at a pre-defined threshold. As a result, LIN codes can simultaneously convey species-level identity (e.g. *C. jejuni* vs *C. coli*), population-level structure (e.g. clonal complexes), and outbreak-level clonal groups - all within a single identifier.

LIN codes require a robust cgMLST scheme as their foundation, so that automated scripts can be used to reliably and efficiently assign LIN codes to new *Campylobacter* genomes in a timely manner. An essential first step to designing the LIN codes in this study was to review the *C. jejuni / C. coli* cgMLST v1 scheme that was published in 2017 by Cody *et al* [22]. We removed problematic loci (inconsistent start sites, internal stop codons, phase variation, paralogs, and frequent assembly or annotation failures) to give the enhanced cgMLST v2 scheme. The cgMLST v2 scheme is more robust than the original scheme, allowing for a lower threshold of 25 *versus* 50 loci for missing data. This is important since missing data can artefactually make isolates appear more related. The cgMLST v2 scheme better captures diversity amongst a global dataset which has greatly expanded since 2017, as well as showing better performance for both species, particularly *C. coli.* While the cgMLST v1 scheme remains available for back-compatibility, the new cgMLST v2 scheme was used to design the LIN codes in this study and is now implemented as the default scheme on PubMLST.

Using a set of 5,664 high-quality, curated genomes and cgMLST v2 profiles, we then defined a series of hierarchical allelic distance thresholds across both *C. jejuni* and *C. coli*, that reflect natural breaks in the population structure and support consistent cluster assignment across time. This is the first published LIN scheme to incorporate two species that are highly recombinogenic, and a similar approach may be needed for other complex bacterial genera. There were additional challenges over previously published single species LIN code systems, however [2, 18, 20]. Whilst the Silhouette and adjusted Wallace scores demonstrated the chosen *Campylobacter* LIN thresholds were cohesive and robust, the adjusted Rand index statistics assessing compatibility with existing MLST nomenclature did not align as well as published schemes for the other bacterial species. This is most likely due to there being two *Campylobacter* species that can recombine but are also genetically diverse, most particularly those from wild birds and environmental waters. In addition, whilst MLST-based CCs are for the most part monophyletic, there is some variation in length of LIN prefix between the CCs, lowering the ARI scores.

The original seven-locus MLST scheme for *Campylobacter* was described nearly 25 years ago based on the sequencing technology available at the time [43]. It has been very successful in identifying species, population structure and host association within *Campylobacter,* given the small number of loci. The cgMLST-based LIN codes give new insight into genetic clustering within *C. jejuni* and *C. coli.* Most isolates start with ‘0’ at the 1^st^ LIN threshold, but four additional species-specific clusters have LIN codes starting with 1-4 and were identified amongst isolates from wild bird and environmental waters. Whilst there may be some debate over whether disparate lineages are distinct species, all except three isolates (n=308) had 100% support for *C. jejuni* or *C. coli* identification by rMLST, yet they split at a higher threshold and are more distinct than the majority of *C. jejuni* and *C. coli* isolates are from each other [23, 44]. In addition, by using LIN codes we can now further investigate clonal complexes that may not be perfectly monophyletic (eg ST-353CC and the *C. coli* ST-828CC), are nested (eg ST-283CC) or are very large, multi-host ‘generalist’ complexes (ST-21CC, ST-45CC and ST-828CC).

Unlike other cgMLST hierarchical clustering algorithms that use single-point identifiers such as HierrCC [45], LIN codes are backwards compatible with existing nomenclature, such as ‘clonal complex’, that are well established in describing the population biology of *Campylobacter*. For example, ‘lineage’ (LIN threshold range 4-6) is equivalent to most CCs and ‘sub-lineage’ (LIN threshold range 7-9) is equivalent to nested CCs or STs (which may overlap in their degree of genetic distinction). ‘Clonal groups’ are equivalent to LIN thresholds 12-17, with between 10 and 1 allele mismatches and describe genetic diversity within STs. Each LIN code integer, up to and including the full LIN profile are searchable amongst isolates and genomes on PubMLST. We have also introduced searchable ‘nicknames’, in line with other organisms such as *Klebsiella pneumoniae [2],* allowing users to search for LIN-defined ST-21CC (with LIN 0_0_0_0_1_0_6 and 0_0_0_0_1_0_8), as well as MLST-defined ST-21CC, for example. The newly designed LIN codes enhance the ability to capture information from WGS data and describe natural breaks in *Campylobacter* population structure that falls across a range of thresholds and cannot be perfectly described by existing nomenclature.

We evaluated the robustness of the chosen LIN thresholds across different applications, including the previously published Havelock North waterborne outbreak in New Zealand [1]. By repeating the analysis of the genomes publicly available from the Havelock North outbreak, our results using LIN codes identified the same clusters of indistinguishable clonal groups, linking human clinical isolates with those from sheep and potable water, implicating agricultural run-off as the source of the outbreak. Our results demonstrated consistency across the two cgMLST schemes, with cgMLST v1 used in the original publication, and cgMLST v2 used in this study. An advantage of using the cgMLST-LIN nomenclature in this application, however, is the ability to immediately identify indistinguishable isolate groups, as well as the LIN thresholds that different clusters are related by.

There are two *C. jejuni* CCs (ST-21CC and ST-45CC) and one *C. coli* CC (ST-828CC) that have been described as large, multi-host ‘generalists’ in terms of their biology [35]. It has been difficult to elucidate their substructure using MLST, and we performed a preliminary investigation using LIN codes to ascertain their usefulness for this application. Using a dataset of 1,800 globally representative, high-quality genomes, ST-21CC could be identified using the 5^th^ LIN threshold, which is at the higher end of the range matching other CCs. LIN thresholds 6-9 successfully distinguished ST-equivalent diversity amongst three prominent STs (ST-19, ST-21 and ST-50) that we investigated in greater detail. Within each of these three STs there was a prominent clonal group, which could be broken down into small groups of 20-30 isolates at the 12^th^ to 17^th^ LIN thresholds, giving clustering levels appropriate for outbreak investigation. In comparison to ST grouping, the LIN codes gave a more even distribution, as well as showing sub-clustering with the ST, and demonstrate promise for furthering understanding transmission within large CCs.

ST-6175 (also grouping with ST-21CC) is gaining attention in the UK, and internationally as a prominent, multi-resistant *C. jejuni* lineage [36]. Indeed, all except one of the isolates from our test global dataset carried the *gyrA* T86I mutation with high predictive correlation for fluoroquinolone resistance[46]. Isolates additionally carried the *tetO* gene, predictive of tetracycline resistance, except for isolates from North America. All isolates within ST-6175 clustered in the 9^th^ LIN threshold, in line with the range shown by other STs. The 11^th^ LIN threshold demonstrated sub-structure within ST-6175, clearly distinguishing clusters by continent and resistance pattern that were stable over time, and even by resistance allele.

The isolate source is imperfectly sampled, with European isolates dominated by human clinical isolates, and North American samples dominated by poultry isolates, which may be a confounding factor. Where human and animal isolates were available together however, they still clustered by continent and resistance pattern, indicating that resistance has expanded by continent, with hints of continued evolution over time. The one cattle isolate sensitive to both fluoroquinolone and tetracycline was from Luxembourg and clustered with other European isolates that were resistant.

Unlike cgMLST-based clustering approaches that rely on dynamic distance cut-offs (e.g. single-linkage thresholds or *ad hoc* grouping by allelic range), LIN codes provide a fixed positional encoding of genomic relatedness. This enables long-term comparability of lineage definitions even as databases grow, a key limitation of previous clustering approaches. As with all cgMLST-based methods, LIN code assignment depends on the quality and completeness of the genome assembly and on the stability of the underlying cgMLST scheme. Differences in gene calling or allele indexing across platforms could affect consistency if not harmonised.

In conclusion, we provide a robust, stable, and publicly accessible LIN code nomenclature for *C. jejuni* and *C. coli* that captures hierarchical genomic diversity and supports high-resolution surveillance. The LIN system is compatible with expanding genome datasets, and new LINs can be assigned independently without requiring reanalysis of the full dataset - a major advantage over distance-based clustering approaches. Future applications of the LIN code scheme include real-time outbreak detection, integration with AMR prediction frameworks, and phylogeographic mapping of transmission pathways. The integration of LIN codes into PubMLST provides a valuable resource for the *Campylobacter* research and public health communities and ensures accessibility for laboratories without specialist bioinformatics support.

## Ethical approval

No ethical approval was required for this study.

## Author contributions

The CReDIT contributor roles taxonomy was used to recognise author contributions as follows. Conceptualization: K.M.P and F.M.C. Methodology: K.M.P, K.A.J., and F.M.C. Software: K.M.P. and K.A.J. Validation: K.M.P., B.P., and F.C. Formal analysis: K.M.P and F.M.C. Investigation: K.M.P., B.P., K.A.J., and F.M.C. Resources: M.C.J.M, K.A.J., S.K.S., and F.M.C. Data curation: K.M.P., K.A.J, and F.M.C. Writing – original draft preparation: K.M.P, B.P., and F.M.C. Writing – review and editing: all authors. Visualisation: K.M.P., B.P., and F.M.C., Supervision: M.C.J.M and F.M.C., Project administration: K.M.P. and F.M.C, Funding acquisition: M.C.J.M, S.K.S. and F.M.C.

## Acknowledgements

The authors would like to acknowledge the use of the Advanced Research Computing (ARC) facilities at the University of Oxford for their computing resources.

## References

1. Gilpin, B.J., et al., A large scale waterborne Campylobacteriosis outbreak, Havelock North, New Zealand. J Infect, 2020. 81(3): p. 390–395.

2. Hennart, M., et al., A Dual Barcoding Approach to Bacterial Strain Nomenclature: Genomic Taxonomy of Klebsiella pneumoniae Strains. Mol Biol Evol, 2022. 39(7).

3. Goddard, M.R., et al., A restatement of the natural science evidence base regarding the source, spread and control of Campylobacter species causing human disease. Proc Biol Sci, 2022. 289(1976): p. 20220400.

4. UKHSA Campylobacter data 2014-2023. 2025.

5. Finsterer, J., *Triggers of Guillain-Barre Syndrome:* Campylobacter jejuni Predominates. Int J Mol Sci, 2022. 23(22).

6. Caron, G., et al., *Campylobacter jejuni Outbreak Linked to Raw Oysters in Rhode Island*, *2021*. J Food Prot, 2023. 86(11): p. 100174.

7. Fernandes, A.M., et al., Partial Failure of Milk Pasteurization as a Risk for the Transmission of Campylobacter From Cattle to Humans. Clin Infect Dis, 2015. 61(6): p. 903–9.

8. Hyllestad, S., et al., *Large waterborne Campylobacter outbreak: use of multiple approaches to investigate contamination of the drinking water supply system, Norway,* June *2019*. Euro Surveill, 2020. 25(35).

9. Kenyon, J., et al., *Campylobacter outbreak associated with raw drinking milk, North West England*, *2016*. Epidemiol Infect, 2020. 148: p. e13.

10. Dolma, K.G., et al., Investigation of an Acute Gastrointestinal Illness Outbreak Linked to Drinking Water in a Higher Educational Institute in East Sikkim, India. Cureus, 2024. 16(7): p. e64050.

11. Jansen, L., et al., *Campylobacteriosis Outbreak Linked to Municipal Water, Nebraska, USA*, *2021(1)*. Emerg Infect Dis, 2024. 30(10): p. 1998–2005.

12. Wensley, A., S. Padfield, and G.J. Hughes, An outbreak of campylobacteriosis at a hotel in England: the ongoing risk due to consumption of chicken liver dishes. Epidemiol Infect, 2020. 148: p. e32.

13. Maiden, M.C., et al., MLST revisited: the gene-by-gene approach to bacterial genomics. Nat Rev Microbiol, 2013. 11(10): p. 728–36.

14. Palma, F., et al., Life Identification Numbers: A bacterial strain approach. 2025: bioRxiv.

15. Vinatzer, B.A., et al., A Proposal for a Genome Similarity-Based Taxonomy for Plant-Pathogenic Bacteria that Is Sufficiently Precise to Reflect Phylogeny, Host Range, and Outbreak Affiliation Applied to Pseudomonas syringae sensu lato as a Proof of Concept. Phytopathology, 2017. 107(1): p. 18–28.

16. Delgado-Blas, J.F., M. Rethoret-Pasty, and S. Brisse, Life Identification Number (LIN) codes for the genomic taxonomy of Corynebacterium diphtheriae strains. Genome Med, 2025.

17. Hadjirin, N.F., et al., Development of a core genome multilocus sequence typing scheme and life identification number code classification system for Staphylococcus aureus. Microb Genom, 2025. 11(8).

18. Jansen van Rensburg, M.J., et al., Development of the Pneumococcal Genome Library, a core genome multilocus sequence typing scheme, and a taxonomic life identification number barcoding system to investigate and define pneumococcal population structure. Microb Genom, 2024. 10(8).

19. Yassine, I., et al., Investigating the population structure of Moraxella catarrhalis using a cgMLST scheme and LIN code system. Nat Commun, 2025. 16(1): p. 9137.

20. Unitt, A., et al. *Neisseria gonorrohoeae LIN codes:* a robust, multi-resolution lineage nomenclature. bioRxiv, 2025.

21. Jolley, K.A., J.E. Bray, and M.C.J. Maiden, Open-access bacterial population genomics: BIGSdb software, the PubMLST.org website and their applications. Wellcome Open Res., 2018. 3: p. 124.

22. Cody, A.J., et al., Core Genome Multilocus Sequence Typing Scheme for Stable, Comparative Analyses of Campylobacter jejuni and C. coli Human Disease Isolates. Journal of Clinical Microbiology, 2017. 55(7): p. 2086–2097.

23. Jolley, K.A., et al., Ribosomal multilocus sequence typing: universal characterization of bacteria from domain to strain. Microbiology, 2012. 158(Pt 4): p. 1005–1015.

24. Wickham H, F.R., Henry L, Müller K, Vaughan D, dplyr: A Grammar of Data Manipulation. 2023: https://github.com/tidyverse/dplyr.

25. Team, R., *RStudio: Integrated Development for R*. RStudio. 2020: PBC, Boston, MA.

26. Wickham, H., Reshaping Data with the reshape Package. Journal of Statistical Software, 2007. 21: p. 1–20.

27. Wickham, H., ggplot2: Elegant Graphics for Data Analysis. 2016: Springer-Verlag New York.

28. Rousseeuw, P.J., Silhouettes: a graphical aid to the interpretation and validation of cluster analysis.. J Comput Appl Math, 1987. 20: p. 53–65.

29. Wallace, D.L., A method for comparing two hierarchical clusterings: comment. J Am Stat Assoc, 1983. 78(383): p. 569–576.

30. Hubert, L.J. and P. Arabie, Comparing partitions. Journal of Classification, 1985. 2: p. 193–218.

31. Scrucca, L., et al., Model-Based Clustering, Classification, and Density Estimation Using mclust in R. 2023: New York, Chapman and Hall/CRC.

32. Jolley, K.A., J.E. Bray, and M.C.J. Maiden, Open-access bacterial population genomics: BIGSdb software, the PubMLST.org website and their applications. Wellcome Open Res, 2018. 3: p. 124.

33. Price, M.N., P.S. Dehal, and A.P. Arkin, FastTree 2--approximately maximum-likelihood trees for large alignments. PLoS One, 2010. 5(3): p. e9490.

34. Didelot, X. and D.J. Wilson, ClonalFrameML: Efficient Inference of Recombination in Whole Bacterial Genomes. PLOS Computational Biology, 2015. 11(2): p. e1004041.

35. Dearlove, B.L., et al., Rapid host switching in generalist Campylobacter strains erodes the signal for tracing human infections. ISME J, 2016. 10(3): p. 721–9.

36. Colles, F.M., et al., Genomics of antimicrobial resistant Campylobacter transmission through UK Agri-Food systems. 2025: Food Standards Agency.

37. Zhou, Z., et al., GrapeTree: visualization of core genomic relationships among 100,000 bacterial pathogens. Genome Res, 2018. 28(9): p. 1395–1404.

38. Sheppard, S.K., et al., Convergence of Campylobacter species: implications for bacterial evolution. Science, 2008. 320(5873): p. 237–9.

39. Oxford, U.o., Enhanced molecular-based surveillance (MLST/whole genome) surveillance and source attribution of Campylobacter infections in the UK. 2019.

40. Colles, F.M., et al., Dynamics of Campylobacter colonization of a natural host, Sturnus vulgaris (European starling). Environmental Microbiology, 2009. 11(1): p. 258–67.

41. Colles, F.M., et al., Comparison of Campylobacter populations in wild geese with those in starlings and free-range poultry on the same farm. Appl and Environ Micobiol, 2008. 74(11): p. 3583–90.

42. Veltcheva, D.L., Re-visiting clonal-complex classifications: a novel machine learning approach for investigating population dynamics of antimicrobial resistance in Campylobacter jejuni. 2023, University of Oxford.

43. Dingle, K.E., et al., Multilocus sequence typing system for Campylobacter jejuni. J Clin Microbiol, 2001. 39(1): p. 14–23.

44. Henaut-Jacobs, S., H. Passarelli-Araujo, and T.M. Venancio, Comparative genomics and phylogenomics of Campylobacter unveil potential novel species and provide insights into niche segregation. Mol Phylogenet Evol, 2023. 184: p. 107786.

45. Zhou, Z., J. Charlesworth, and M. Achtman, HierCC: a multi-level clustering scheme for population assignments based on core genome MLST. Bioinformatics, 2021. 37(20): p. 3645–3646.

46. Aleksic, E., et al., Resistance to Antibiotics in Thermophilic Campylobacters. Front Med (Lausanne), 2021. 8: p. 763434.

