## Supplemental file 1 for "A stable, hierarchical LIN code system for *Campylobacter jejuni* and *Campylobacter coli:* A unified genomic nomenclature for lineage-level typing and global surveillance"

**Supplementary figures**

**Figure S1.** Genome statistics (A) assembly size, (A) contig count, (B) N50 and (C) GC content, shown for (a) development dataset 1, and (b) development data set 2 (n=1,781 isolates).

**(a) Development dataset 1 (n=5,664 isolates)**

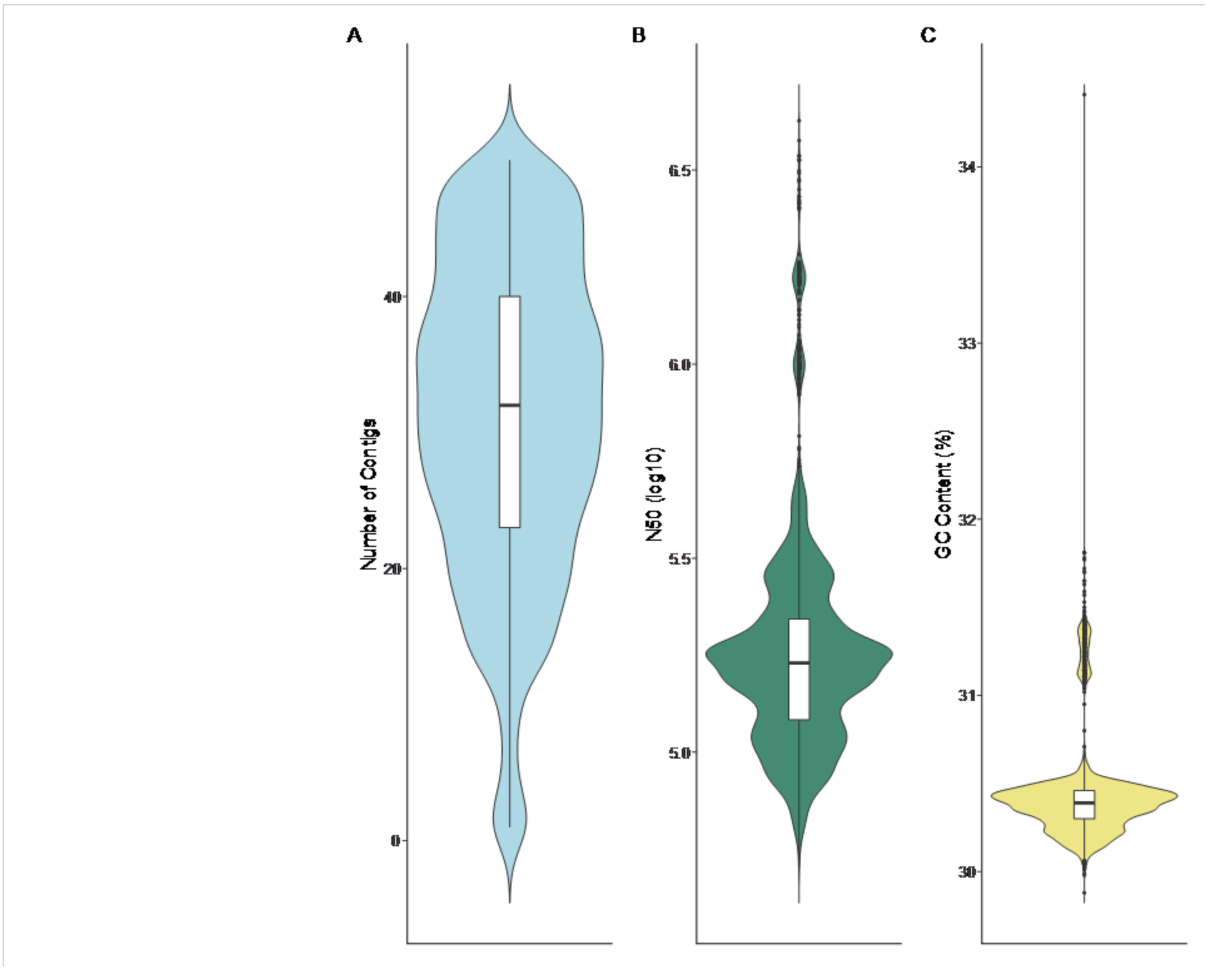


**(b) Development dataset 2 (n=1,781 isolates)**

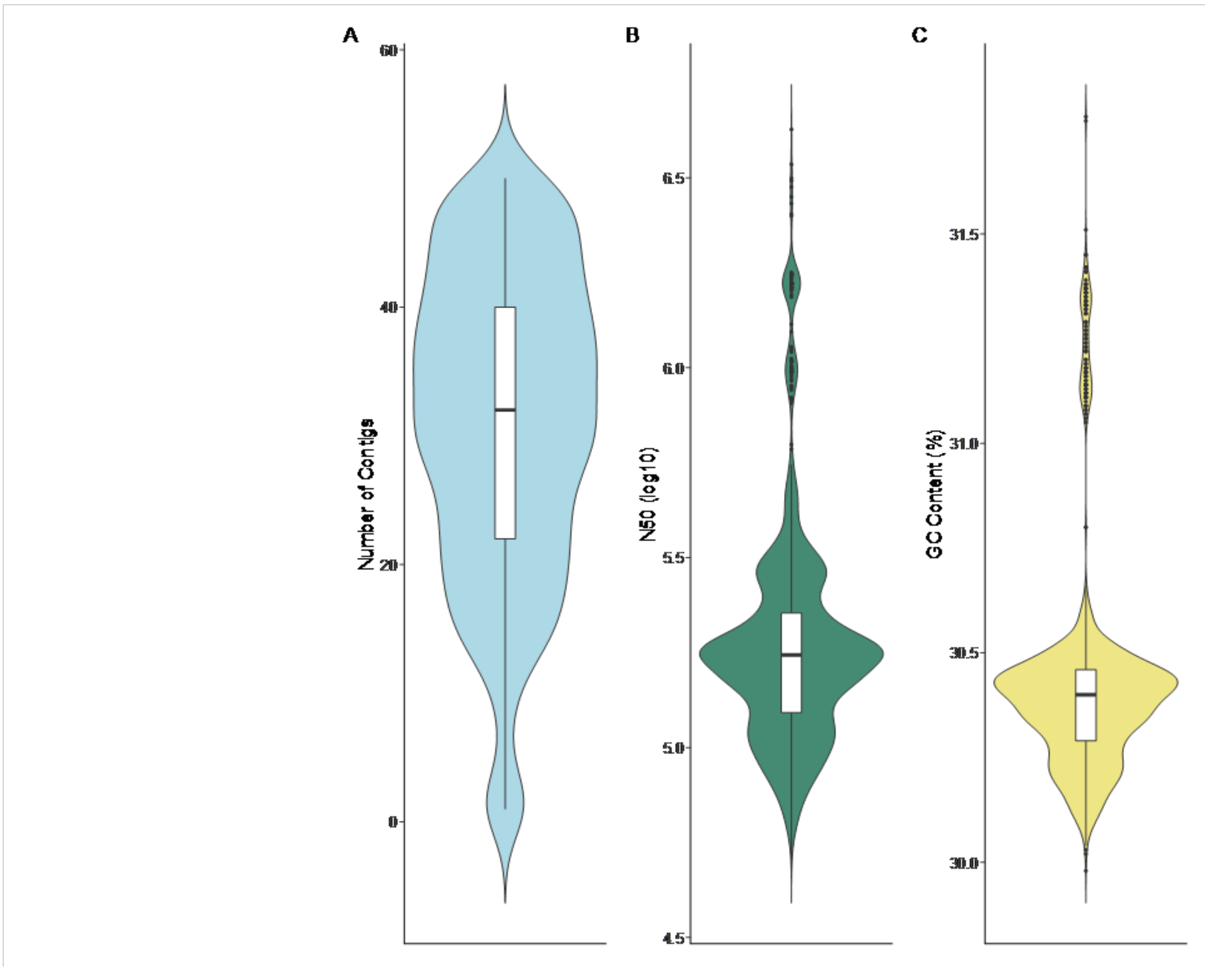


**Figure S2. Clustering of ST-45CC, ST-283CC and ST-179CC isolates, shown by (a) CC, and (b) LIN code threshold-3, using Grapetree analysis.**

(a)

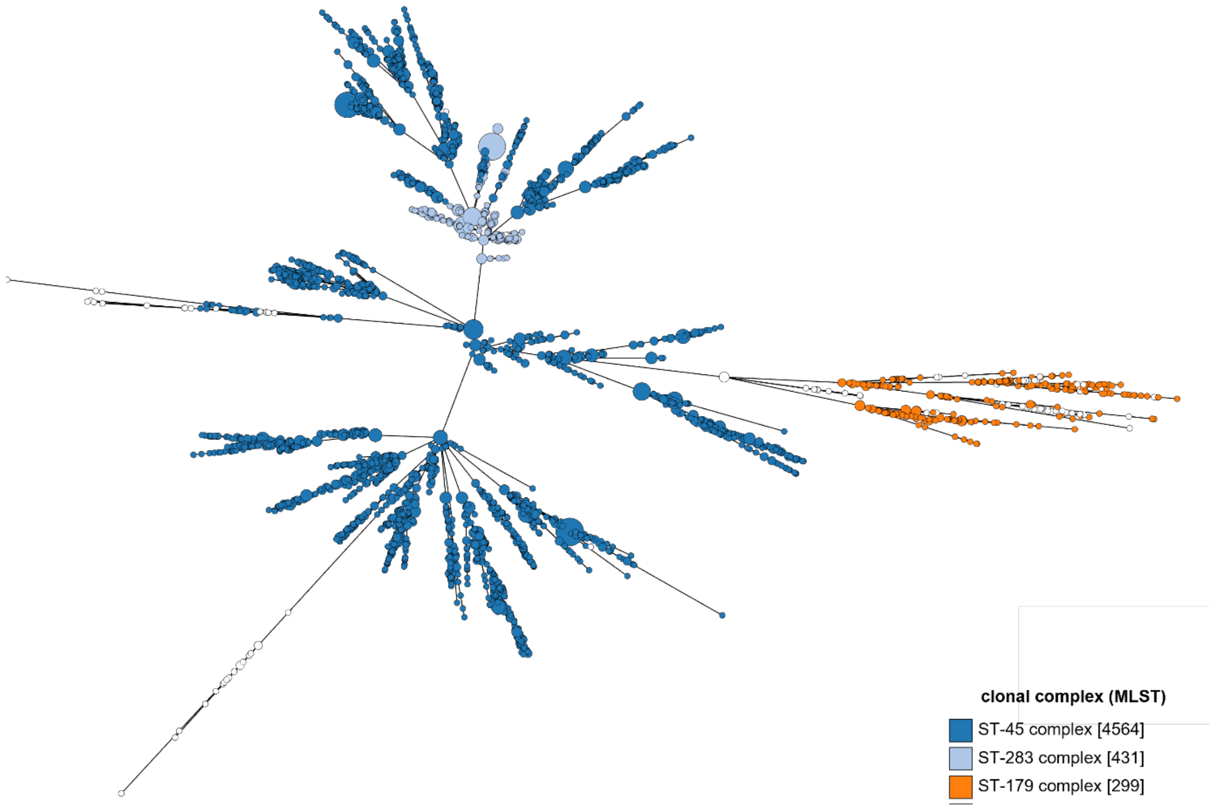
clonal complex (MLST)
ST-45 complex [4564]
ST-283 complex [431]
ST-179 complex [299]

(b)

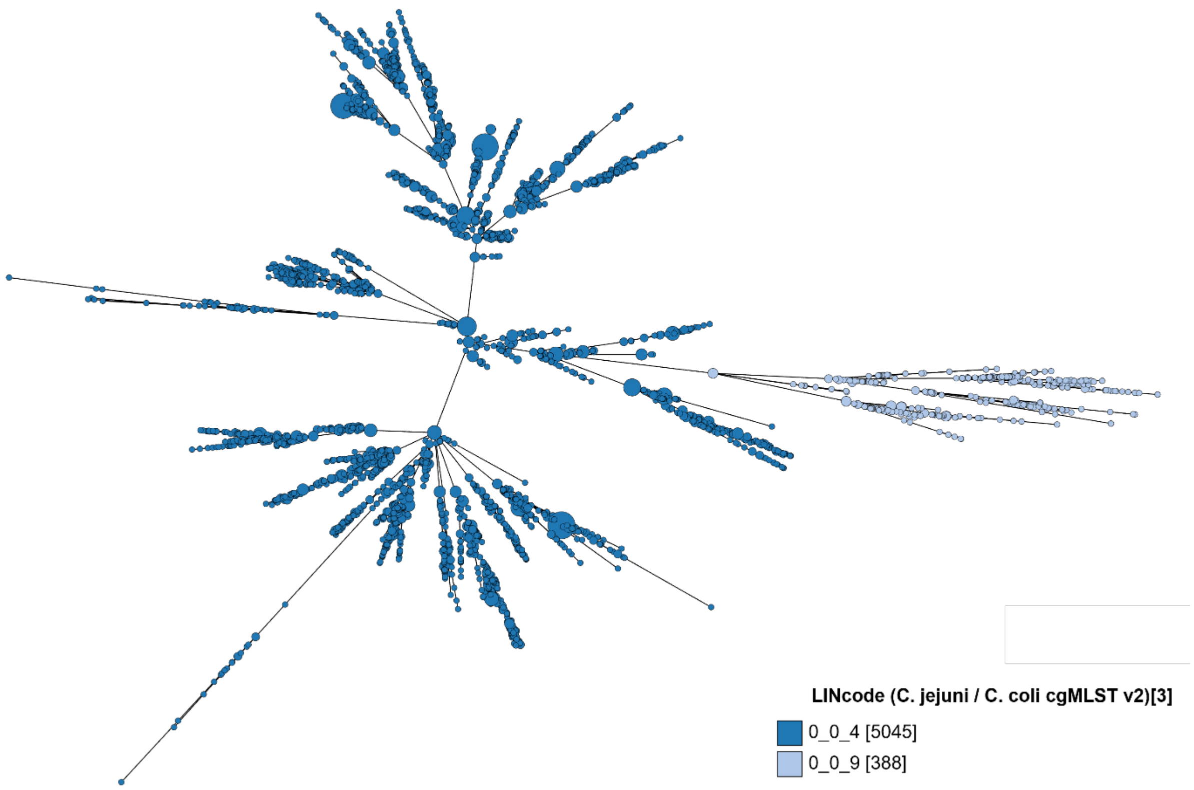
LINcode (C. jejuni / C. coli cgMLST v2)[3]
0_0_4 [5045]
0_0_9 [388]

**Figure S3.** The distribution of isolates from the ST-21CC test dataset showing distribution by (a) ST and (b) LIN code at threshold-9.

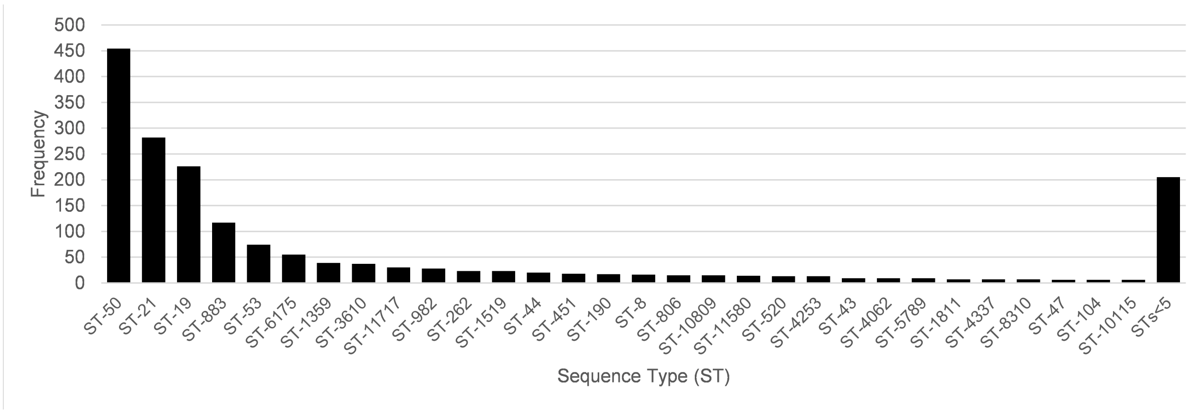


(b)

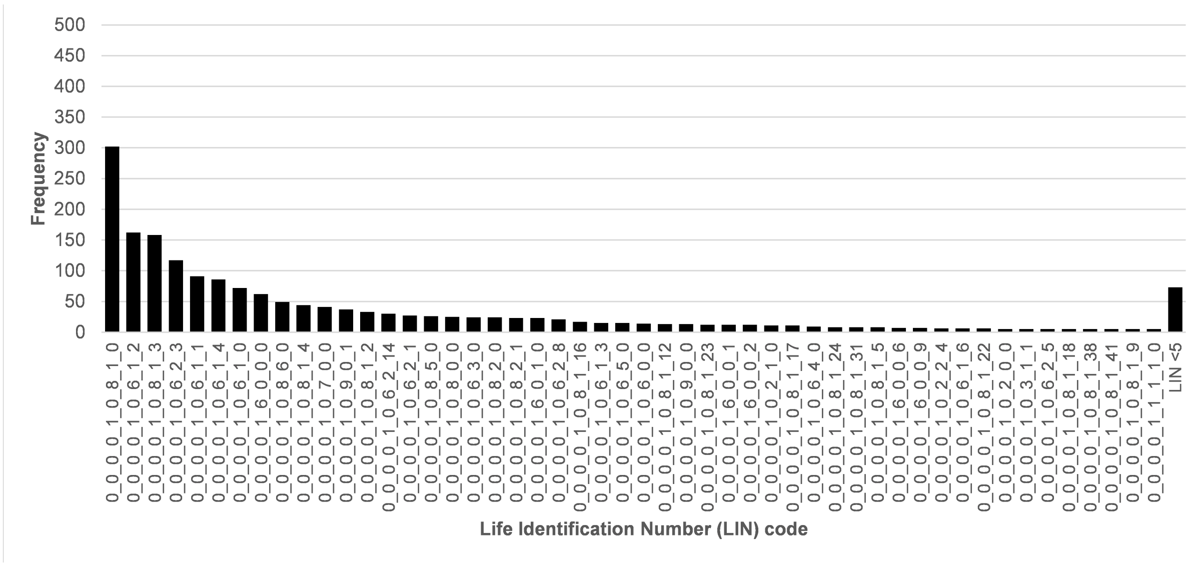


**Figure S4.** The clustering of isolates within ST-6175 (ST-21CC) using the cgMLST v2 scheme and labelled by (a) LIN threshold-9, (b) *gyrA* allele (CAMP0950) predictive for fluoroquinolone resistance and (c) *tetO* allele (CAMP1698) predictive of tetracycline resistance.

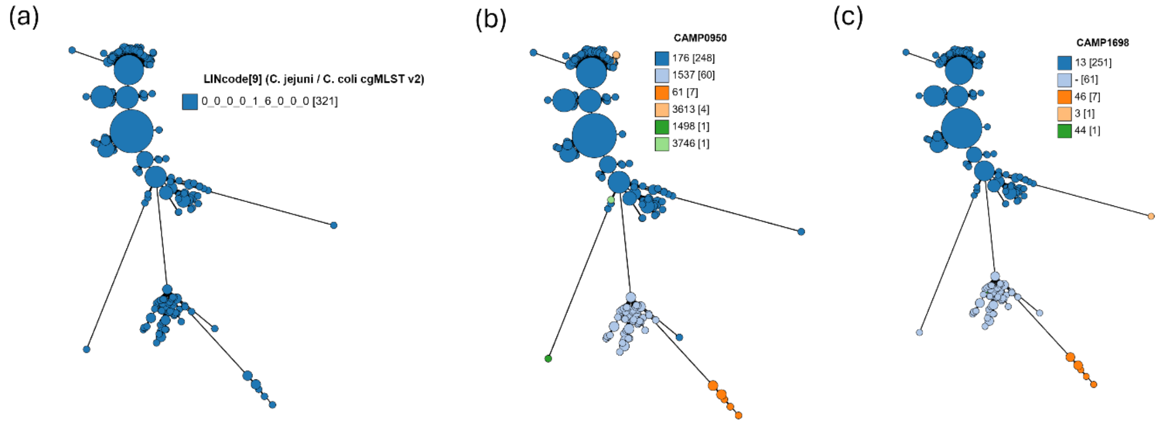


**Figure S5.** The clustering of isolates within ST-828 using the cgMLST v2 scheme and labelled by (a) CC, (b) ST, (c) LIN code Threshold-4, (d) LIN code Threshold-6, (e) LIN code Threshold-7, (f) LIN code Threshold-8, shown using GrapeTree.

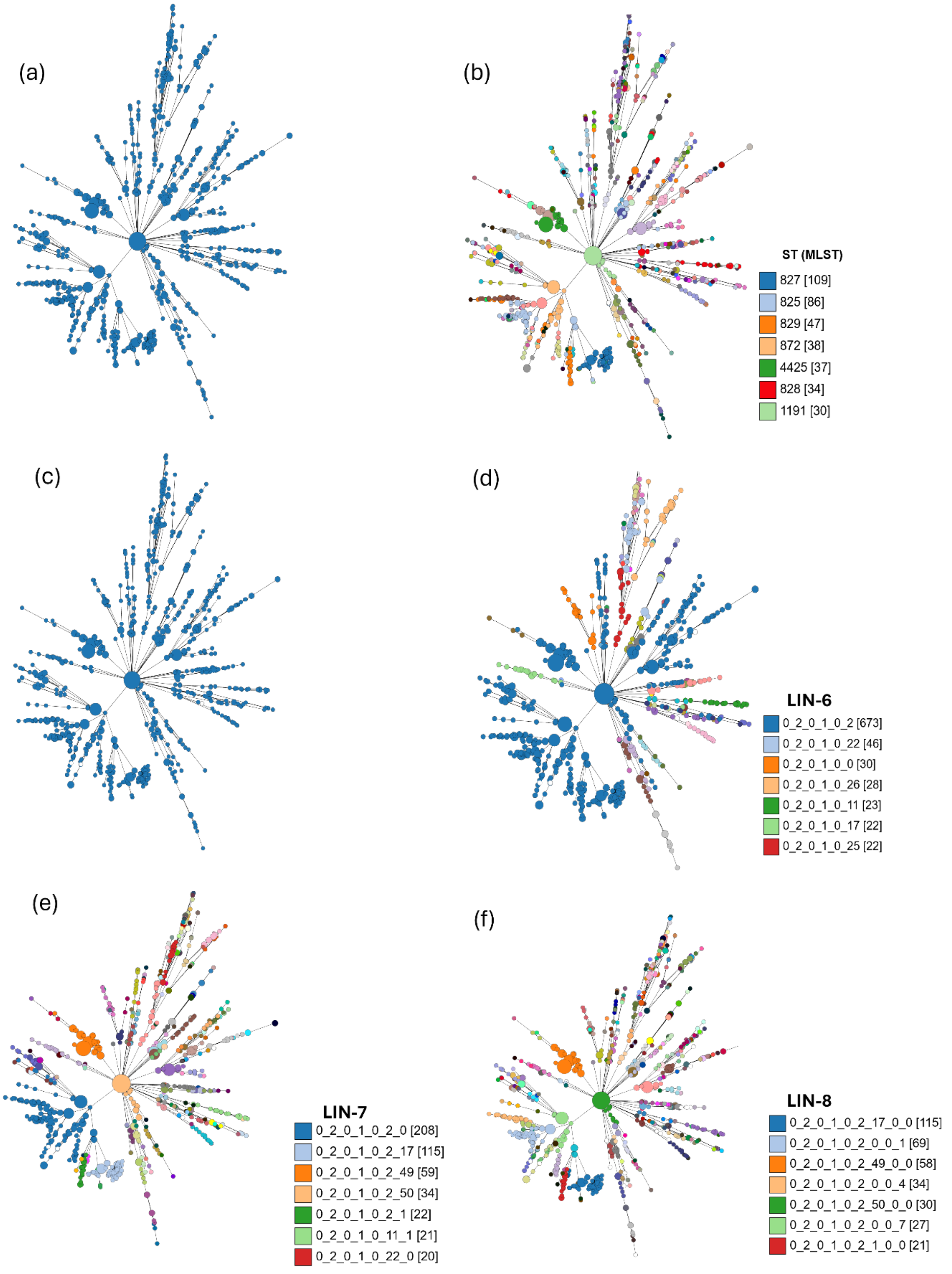


**Figure S6.** The clustering of isolates within ST-828 (ST-828CC) using the cgMLST v2 scheme and labelled by (a) ST-828, (b) LIN code Threshold-6, (c) continent, (d) source, shown using GrapeTree.

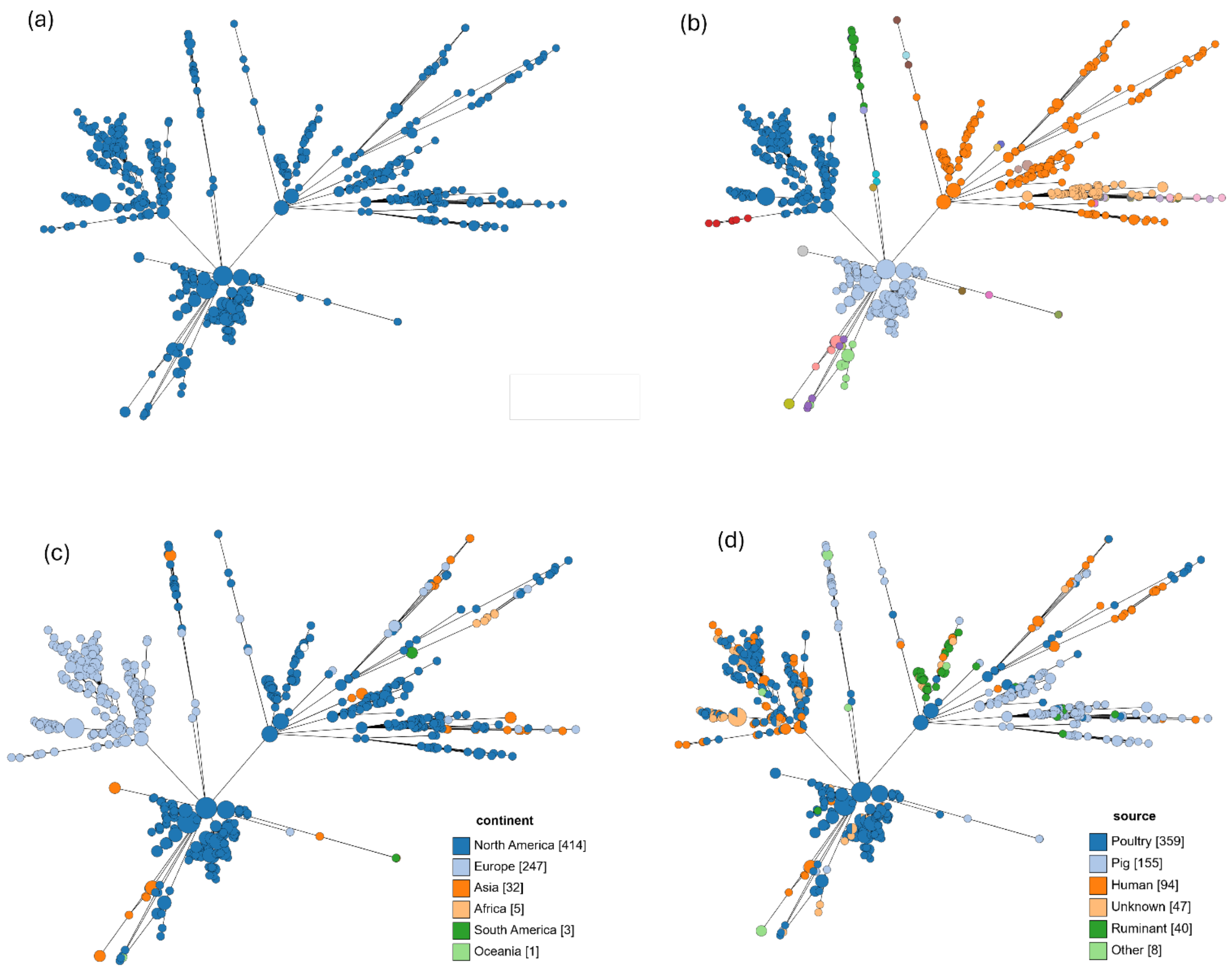
(a)
(b)
(c)
(d)
continent
North America [414]
Europe [247]
Asia [32]
Africa [5]
South America [3]
Oceania [1]
source
Poultry [359]
Pig [155]
Human [94]
Unknown [47]
Ruminant [40]
Other [8]

**Table S2.** LIN code thresholds that distinguish genomes based upon MLST clonal complex (CC) 'nickname' nomenclature. They are searchable on PubMLST using clonal complex (*C. jejuni I C. coli* cgMLST v2), distinct from clonal complex (MLST). Note, CCs 1264, 1304, 1325, 1347 and 443 contained between 2 and 20 genomes, and no cgSTs or LIN codes were available.

| Species | Clonal Complex | LIN prefix |
|---|---|---|
| *Campylobacter jejuni* | ST-21 complex | 0_0_0_0_1_0_6 and 0_0_0_0_1_0_8 |
| | ST-22 complex | 0_0_0_0_1_23 |
| | ST-41 complex | 0_0_0_0_2_0 |
| | ST-42 complex | 0_0_0_0_1_22 |
| | ST-45 complex | 0_0_4_0 (could not resolve from ST-283CC) |
| | ST-48 complex | 0_0_0_0_1_0_2 |
| | ST-49 complex | 0_0_0_0_1_17 |
| | ST-52 complex | 0_0_0_0_17 |
| | ST-61 complex | 0_0_0_0_1_0_0 |
| | ST-177 complex | 0_0_10_0_0_0 |
| | ST-179 complex | 0_0_9_0 |
| | ST-206 complex | 0_0_0_0_1_0_3 and 0_0_0_0_1_0_4 |
| | ST-257 complex | 0_0_0_0_11 |
| | ST-283 complex | 0_0_4_0 (could not resolve from ST-45CC) |
| | ST-353 complex | 0_0_0_0_1_9 |
| | ST-354 complex | 0_0_0_0_1_18 |
| | ST-362 complex | 0_0_0_0_2_2 |
| | ST-403 complex | 0_0_0_0_1_19 |
| | ST-443 complex | 0_0_0_1_0_0_0 |
| | ST-446 complex | 0_0_0_0_4 |
| | ST-460 complex | 0_0_0_0_9 |
| | ST-464 complex | 0_0_0_0_1_13 |
| | ST-508 complex | 0_0_8_0 |
| | ST-573 complex | 0_0_0_4_1 |
| | ST-574 complex | 0_0_0_0_1_26 |
| | ST-581 complex | 0_0_0_18 |
| | ST-607 complex | 0_0_0_0_1_27 |
| | ST-658 complex | 0_0_0_0_0 |
| | ST-661 complex | 0_0_0_4_0 |
| | ST-677 complex | 0_0_19_0 |
| | ST-682 complex | 0_0_10_0_0_1 |
| | ST-692 complex | 0_0_0_4_4_1 |
| | ST-702 complex | 0_0_0_4_4_3 |
| | ST-952 complex | 0_0_16 |
| | ST-1034 complex | 0_0_0_4_7 |
| | ST-1275 complex | 0_0_11_0 |
| | ST-1287 complex | 0_0_0_8 and 0_0_0_9 |
| | ST-1332 complex | 0_0_0_4_4_0 |
| *Campylobacter coli* | ST-828 complex | 0_2_0_1 |
| | ST-1150 complex | 0_2_0_0 |
